# Histone acetylation insulator SET orchestrates PP2A inhibition and super-enhancer activation

**DOI:** 10.1101/2023.01.15.524091

**Authors:** He Xu, Di Wu, Jin Xu, Yubin Lei, Yalan Lei, Xianjun Yu, Si Shi

## Abstract

Wide-spread growth-essential genes are hyper-transcribed in the pancreatic cancer cells. Searching for the factors that reprogram this abnormal transcription, we identified the nuclear oncogene SET that supported CDK9-induced and Pol II-mediated transcription. SET disrupted PP2A-A/C interaction via its C-terminal domains. Through blocking PP2A activity, SET assisted CDK9 to maintain Pol II CTD phosphorylation and activated mRNA transcription. Meanwhile, as a histone acetylation insulator, SET mainly suppressed histone acetylation in the gene promoters but evaded enhancers. Massive super-enhancer associated genes, including the oncogene *MET*, were hence permitted to be transcribed by SET over-expression. Our findings position SET as a key factor that bridges histone acetylation and PP2A related transcription in cancer cells.

## Introduction

Non-mutational epigenetic reprogramming is increasingly discussed as a driving force in tumorigenesis (Baylin and Jones, 2016; Flavahan et al., 2017; Hanahan, 2022). Several lines of evidence have indicated the existence of global enhancement of transcription that favors the cancer growth. For instance, c-Myc up-regulation has long been identified as a cancer “driver” (Meyer and Penn, 2008; Nesbit et al., 1999) and c-Myc was found to amplify almost all the mRNA-transcribing genes in cancer cells (Lin et al., 2012). For another example, aberrant activation of super-enhancers, which associated with the hyper-transcription of massive critical oncogenes, was observed in multiple cancers (Chapuy et al., 2013; Chipumuro et al., 2014; Christensen et al., 2014; Hnisz et al., 2013; Kwiatkowski et al., 2014; Loven et al., 2013; Mansour et al., 2014; Sengupta and George, 2017; Thandapani, 2019).

Moreover, histone acetylation reader BRD4 recruits Cyclin T/CDK9 complex to phosphorylate Pol II carboxy terminal domain (CTD) and hence serves as a genome-wide activator of mRNA transcription (Allen and Taatjes, 2015; Jonkers and Lis, 2015; Lu et al., 2015; Peterlin and Price, 2006; Zhou et al., 2012). And intriguingly, BET inhibitors that specifically target BRD4 have shown to be selective anti-proliferation compounds in the pre-clinical treatment of leukemia and solid tumors (Filippakopoulos et al., 2010; Loven et al., 2013; Roe et al., 2015; Yan et al., 2019). In the opposite, PP2A, a tumor suppressor, counteracts with the BRD4-CDK9 axis to dephosphorylate Pol II CTD and attenuates global hyper-transcription (Tellier et al., 2022; Vervoort et al., 2021; Zheng et al., 2020).

Global hyper-transcription reflects the epigenetic plasticity of the cancer cells and it has been supposed that chromatin status permissive for transcription onset would favor the cancer cells sampling for the fitness genes to promote cell growth (Flavahan et al., 2017). However, the epigenetic factors that switch the chromatin status to prompt the global hyper-transcription remain largely elusive.

Nuclear oncogene SET (also called TAF-I) was first described as a nuclear factor that stimulated transcription from chromatin template (Gamble et al., 2005; Matsumoto et al., 1995; Okuwaki and Nagata, 1998). But later it was also purified as a component of INHAT (inhibitor of histone acetyltransferases) complex that negatively regulated histone acetylation (Seo et al., 2001) and also a potent inhibitory protein of PP2A (Li et al., 1996; Neviani et al., 2005; Saito et al., 1999; Trotta et al., 2007). Later it was found that the acid tail of SET could bind the lysine-rich domain of histones or other proteins to prevent them from being acetylated. So, SET is an “insulator” of protein acetylation (Chae et al., 2014; Wang et al., 2016). The multi-task nature of SET molecule led to the very controversial conclusions regarding its function (Asai et al., 2020; Edupuganti et al., 2017; Kalousi et al., 2015; Krishnan et al., 2017). SET was reported to remodel the chromatin structure and facilitated transcription (Fan et al., 2003; Gamble et al., 2005; Haruki et al., 2006; Kato et al., 2007; Miyaji-Yamaguchi et al., 1999; Okuwaki and Nagata, 1998). And SET was also found to integrating chromatin hypoacetylation and transcriptional repression (Cervoni et al., 2002; Karetsou et al., 2004; Matsumoto et al., 1999).

In search for the driver factors related to the hyper-transcription in the pancreatic cancer, we found that SET actually supported CDK9-induced and Pol II-mediated transcription, which was related to the function of SET as a PP2A-A/C disruptor. SET did suppress histone acetylation mainly in the gene promoters but not in the enhancers, which permitted the hyper-acetylation of super-enhancers in concert with SET-induced hyper-transcription of the massive oncogenes including *MET*.

## Results

### The massive hyper-transcription of growth-essential genes in pancreatic cancer

To investigate the changes of transcriptome landscape in pancreatic cancer, we explored the differently expressed genes (DEGs) in pancreatic duct adenocarcinoma (PDAC) compared with normal pancreas tissue. We expected the number of the up-regulated genes and the down-regulated genes would be roughly even. However, for whatever the cut-off criteria we chose, significantly more up-regulated genes were observed in the PDAC than the down-regulated (**Figure 1A**). Considering the possibility that normal tissue in the TCGA database might hold some bias in the gene transcription level, we further looked into the DEGs in multiple PDAC cell lines compared with other cancer cell lines. Still, the result showed that PDAC cells preferred to elevate gene transcription level rather than to turn them down (**Figure S1A**). We were curious whether the massive up-regulated genes in the PDAC were randomly selected by the cancer cells or elaborately chosen based on their cellular function. So, we calculated the average CERES values of those DEGs in 44 types of PDAC cells using the DepMap data (Meyers et al., 2017), and compared the CERES scores of the up-regulated genes with the down-regulated ones. We found the up-regulated genes had statistically lower CERES values (**Figure S1B**), which means the up-regulated genes in the PDAC are prone to be more essential for cancer cell growth (Meyers et al., 2017) while the down-regulated genes not. Taking these together, a transcription remodeling program favoring the cancer growth is obvious in PDAC.

**Figure 1.**
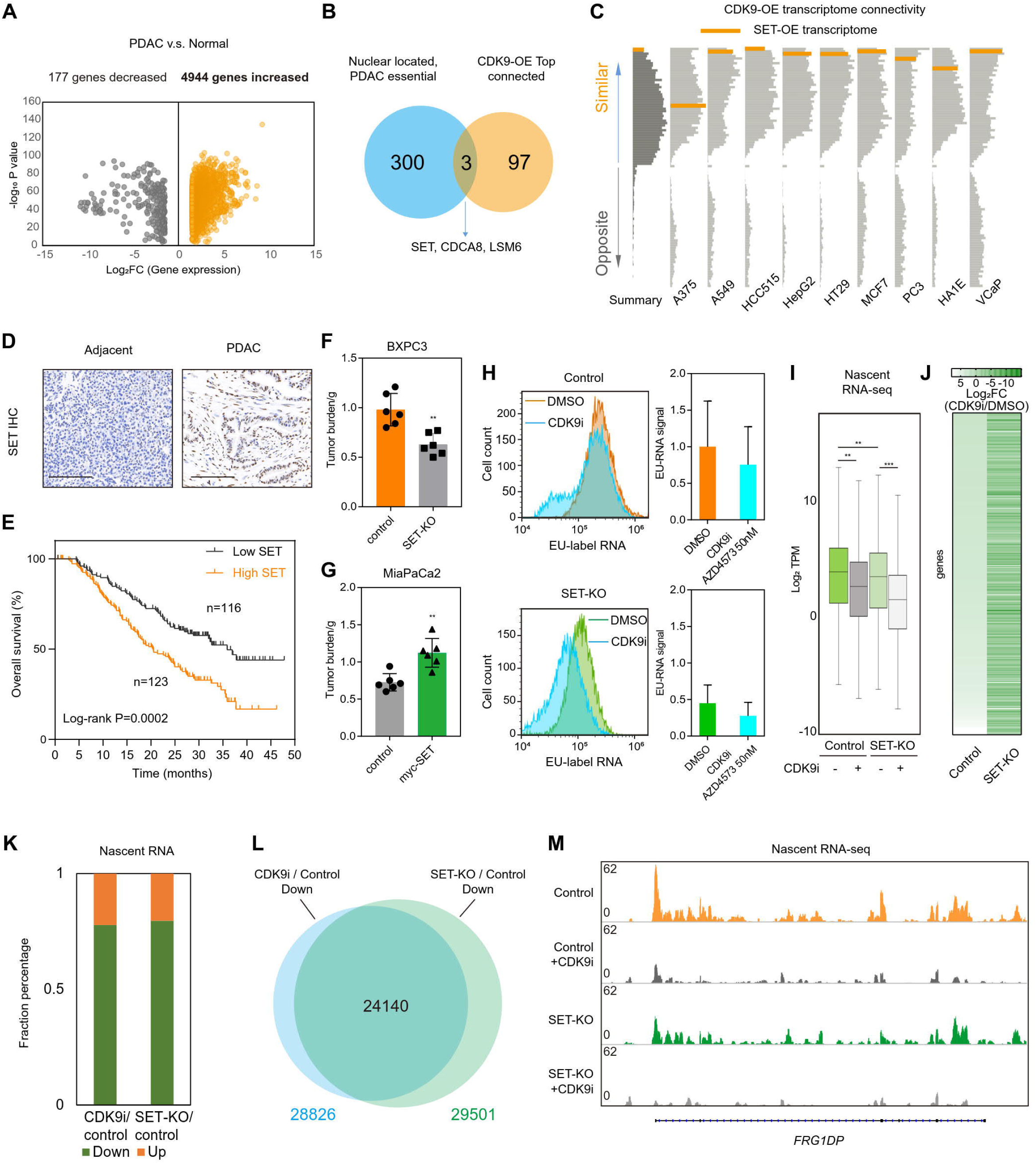
The oncogene SET supports CDK9-induced high level of transcription in cancer cells. (A) Scatter plot showing the differently expressed genes (DEGs) in Pancreatic Duct Adenocarcinoma (PDAC) compared with normal pancreas tissue, Log_2_FC>1.5 or <-1.5, P<0.01; based on the TCGA data. (B) Venn diagrams indicate 303 PDAC essential genes that encode nuclear proteins (see also **Figure S1C**) and top 100 genes that induced the most similar transcriptome signature like CDK9 over-expression in multiple cancer cell lines. (C) Cell connectivity map (L1000 platform) showing that SET-OE induced most similar transcriptome signature like CDK9-OE in 9 different cancer cell lines. (D) Immunohistochemistry (IHC) using SET-specific antibody in PDAC tissue sections. Scale bar: 200μM. (E) Based on the SET IHC scores, a cohort of 278 PDAC patients was divided into SET high and SET low groups, the survival plot was drawn. (F) BXPC3 control cells or SET-KO cells, and (G) MiaCaPa2 control cells or myc-SET stably overexpressing cells were subcutaneously implanted into the nude mice for 28 days before tumor burden was measured. (H) BXPC3 control cells or SET-KO cells (treated w/o CDK9 inhibitor AZD4573 for 2 hrs) were pulse-labeled with EU for 30 min and the nascent RNA incorporated with EU was further covalently labeled with fluorescent via Click chemistry, the labeled cells were analyzed with FACS. (I) Box plot shows the overall changes of nascent RNA in BXPC3 control or SET-KO cells (treated w/o AZD4573). (J) Heatmap shows the gene nascent transcripts fold change (treated w/o AZD4573) in BXPC3 control or SET-KO cells, the top down-regulated genes in the control cells were shown. (K) Fractional bar chart indicates the portion of down/up regulated nascent RNA in the indicated treatments. (L) Venn diagram indicates the overlapping of down regulated nascent RNA either after CDK9 inhibitor treatment or in SET-KO cells. (M) Tracks profile the nascent RNA expression in a representative gene locus.

### The nuclear oncogene SET, elevated in PDAC, is essential for PDAC cells and supports CDK9-induced hyper-transcription

To find out the mechanism underlying the transcription reprogramming in PDAC cells, we planned to screen for genes that drive the abnormal elevation of transcription. Rather than the genes impeding general transcription, it is more difficult to screen out the transcription-promoting genes using the EU-labeled RNA-FACS system because knock-out of the target genes as well as the cell-essential genes would equally turn off the EU-label signals. Alternatively, we focused on the genes that had been elevated in the PDAC. We conjectured, those driver genes to induce hyper-transcription were also in this subset, so they must be (a) PDAC-essential while not common-essential; (b) encoding nuclear proteins related to transcription process. Applying this filtering standard, we finally got 303 candidate genes of which 93 genes are known to be closely related to transcription (**Figures 1B** and **S1C**).

Among the different functional categories of these genes, we were interested with one named “Chromatin modifier complex” because it is most enriched with histone acetylation modifiers (**Figure S1D**). In PDAC, these up-regulated genes were fairly correlated with each other in the mRNA levels (**Figure S1E**), implying their functional relevance. Since histone acetylation is strongly related with gene expression in eukaryotic cells, this small subset of genes seemed worth deeper investigation.

In another dimension, we focused on the clue that BRD4-CDK9 axis plays the pivotal role in the histone acetylation-related transcription activation (Jonkers and Lis, 2015; Peterlin and Price, 2006). We speculated, those genes driving the global hyper-transcription were possibly related to the CDK9 axis. So up-regulation of those genes would render similar transcriptome changes like CDK9 gain-of-function. Based on this speculation, we searched for the genes that induced the most similar transcriptome changes like CDK9 over-expression (**Figure S1F**) using a transcriptome matching method (Lamb et al., 2006; Subramanian et al., 2017). When we overlapped the top 100 CDK9-OE connected genes with the 303 candidate genes mentioned above, we found only 3 genes (that are up-regulated in PDAC, essential for PDAC, encoding nuclear protein, and related with CDK9 functional axis, **Figure 1B**). In the 3 genes, we found that over-expression of nuclear oncogene SET induced the most similar transcriptome phenotype like CDK9 gain-of function (**Figures 1C** and **S1F**), which strongly implied the function of SET gene in the positive regulation of mRNA transcription. Moreover, the SET gene is also one of the members of the histone acetylation modifier complexes noted above (**Figure S1D**). All these hints made SET gene a very special candidate that might be involved in the wide-spread hyper-transcription in PDAC.

SET was reported to be a putative nuclear oncoprotein (Adler et al., 1997; Christensen et al., 2011). In order to verify the cancer-related function of the SET gene, firstly we quantified the mRNA level of SET in a panel of 33 types of cancers. SET mRNA is substantially up-regulated in 11 cancer types including the pancreatic cancer (**Figure S1G**). IHC staining using SET specific antibody revealed increasing nuclear signal in PDAC tissues other than the adjacent non-cancer tissues (**Figures 1D** and **S1H**).

Patient cohort study verified that higher SET mRNA or protein levels predicted poor outcomes and shorter survival time (**Figures 1E** and **S1I**). Via CRISPR-Cas9 technology and stable transfection, we constructed SET loss-of-function and gain-of-function lines in different PDAC cells (**Figure S1J**). Xenograft experiments showed less tumor-burden of SET-KO PDAC cells (**Figure 1F**) but contrary phenotype in the SET-OE cells (**Figure 1G**). In vitro studies also demonstrated similar function of the SET gene (**Figures S1K** and **S1L**).

To testify whether SET gene is associated with (CDK9 induced) hyper-transcription, we measured the gross transcription rate in the SET-OE or SET-KO cells by quantifying EU-labeled nascent RNA after EU pulse labeling. Comparing with the control cells, SET-OE cells transcribed more nascent RNA (**Figure S1M**), while SET-KO cells transcribed significantly less RNA in a given time (**Figure 1H**). In the control cells, CDK9 inhibitor also decreased nascent RNA amount. But in SET-KO cells, the same concentration of CDK9 inhibitor decreased nascent RNA amount to a lower level (**Figure 1H**). More precise sequencing of the nascent RNA indicated that SET-KO decreased the global nascent mRNA level synthetically with CDK9 inhibition (**Figure 1I**). Detailed study of the nascent RNA-seq showed the major part of the mRNA, which were down-regulated by CDK9 inhibitor in the control cells, would be decreased to a much lower level in the SET-KO cells (**Figures 1J** and **S1N**). Since either CDK9 inhibitor or SET-KO would mainly suppress nascent mRNA transcription (**Figure 1K**), and the bulk of the down-regulated nascent mRNAs either by CDK9 inhibition or by SET-KO were largely overlapped (**Figure 1L**), it echoes the phenotype previous observed that CDK9-OE and SET-OE induced most similar transcriptome changes (**Figures 1C** and **S1F**). Our discoveries indicate the role of SET gene in supporting CDK9-induced transcription, as exemplified in **Figure 1M**.

### Loss of SET decreased Pol II engagement at target genes, reducing the mRNA transcription of these genes

Next, we analyzed the effect of SET-KO on the stable mRNA level through regular RNA-seq. We carried out GO analysis of the down-regulated genes after SET knock-out and found these genes were the common targets of transcription regulatory factors and Pol II RNA polymerase (**Figure 2A**). These genes constituted a functional network in turn to exert gene transcription and RNA metabolism (**Figure 2B**). These observations linked the function of SET gene to Pol II and general transcription machinery. To explore more information about SET target genes, we performed SET CUT&Tag to identify the genome-wide SET binding sites. Also, we grouped the DEGs after SET-KO into different subsets according to the mRNA fold changes. We found the down-regulated genes after SET-KO had bound more SET in the control cells (**Figure 2C**). It means the genes directly targeted by SET are inclined to be turned down after SET-KO (**Figures 2D** and **S2A**), which further validated SET gene’s function of promoting transcription. Similarly, the up-regulated genes after SET-OE also had more SET binding than the down-regulated genes (**Figure S2B**). It is noteworthy that the most down-regulated genes after SET knock-out had lower CERES scores than the up-regulated, which implies that SET prefers to support growth-essential genes to transcribe (**Figure S1C**).

**Figure 2.**
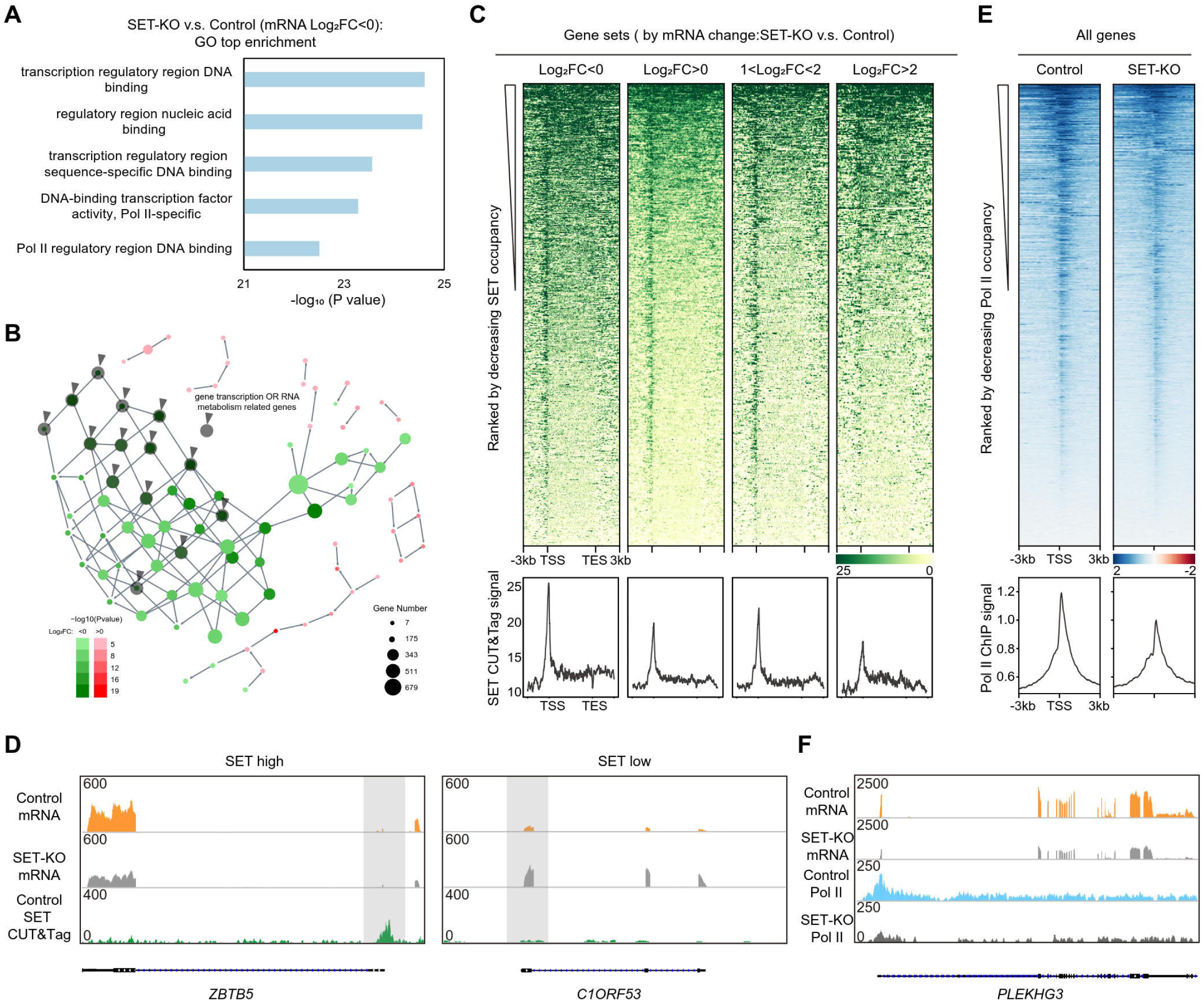
Loss of SET decreased Pol II engagement at target genes, reducing the mRNA transcription of these genes. (A) mRNA-seq defines the down-regulated genes after SET knock-out, the diagram shows the GO analysis result of these decreasing genes. (B) Directed network illustrates the functional clustering of the decreasing genes after SET loss. (C) Heatmaps and metaplots of SET CUT&Tag signals on indicated genes ranked by decreasing occupancy. (D) Tracks profile the mRNA expression and SET occupancy at representative gene loci. (E) Heatmaps and metaplots show the Pol II ChIP-seq signals at scaled TSS vicinity regions ranked by decreasing occupancy. (F) Tracks profile the mRNA expression and Pol II occupancy at a representative gene locus.

To compare SET gene’s effects on nascent RNA transcription and stable mRNA level. We found the genes with both decreased nascent RNA and down-regulated mRNA (**Figure S2D**). Again, GO analysis indicated these genes were the common targets of general transcription factors and Pol II (**Figure S2D**). This fact led us to analyze the function of Pol II in the control and SET-KO cells.

Pol II ChIP-seq analysis showed, comparing with those in the control cells, both genome-wide engagement of Pol II and promoter-specific binding of Pol II were significantly decreased in the SET-KO cells (**Figures 2E**, **2F** and **S2E**). However, calculating the Pol II pausing index (indicated as Pol II Traveling Ration, Pol II TR) in the control and SET-KO cells, as well as in different subset of genes, we found no differences (**Figure S2F**). We speculated that SET-KO had induced a substantial decrease of available Pol II engaged to the chromatin, at least preventing the transcription initiation steps.

Since SET had shown to be able to prompt massive hyper-transcription, we would like to know whether the up-regulated genes in PDAC were related to SET gain-of-function. We identified 574 core up-regulated genes in PDAC (**Figure S2G**) and observed that 57.8% in the mRNA (**Figure S2H**) and 77.2% in the nascent RNA of these genes were turned down by SET-KO (**Figure S2I**). These results demonstrated that SET is one of the driver genes that enhanced the global transcription in PDAC.

### SET inhibits PP2A activity via disrupting PP2A-A/C complex

In order to elucidate the mechanism of SET gene function in supporting both CDK9-induced and Pol II-mediated transcription, we analyzed the SET bound proteins in the control and SET-OE cells using mass spectrum. In the control cells, SET bound to multiple factors involved in RNA metabolism and gene expression (**Figure S3A**). But in the SET-OE cells, only a part of these factors were more enriched by excessive SET protein (**Figure 3A**). We hence identified the specific binding partners of SET by comparing SET interactomes in the control and SET-OE cells. Apart from the known binding partners of SET, for example ANP32A (PP32) (Seo et al., 2001), we found PP2A catalytic subunit (PPP2CA or PPP2CB) is another specific binding partner of SET (**Figures 3A** and **S3B**). In contrast, PP2A-A, PP2A-B or non-canonical PP2A-B factors including the INTAC complex components were almost all depleted in the SET-OE cells (**Figures 3A** and **S3B**). In fact, considering the distinct situations in different cell lines as we analyzed, PP2A-C and histone H3 were the only specific binding targets of SET (which could be more enriched by SET in the SET-OE cells versus the control cells, **Figure S3C**). It has been discovered that PP2A inhibits transcription via antagonizing CDK9 phosphorylation of Pol II CTD domain (Vervoort et al., 2021; Zheng et al., 2020). It seemed possible that SET gene function in supporting CDK9-induced and Pol II-mediated transcription is likely related to the PP2A. Using co-IP and western blot experiments, we verified the interaction between SET and PP2A-C proteins (**Figure 3B**). SET has been purified as an endogenous inhibitory protein of PP2A activity (Li et al., 1996; Neviani et al., 2005). We also observed that PP2A activity was significantly suppressed in the SET-OE cells (**Figure 3C**). Intriguingly, also in the SET-OE cells, the interaction between PP2A-C and PP2A-A/B was impaired (**Figure 3D**). Therefore, we conjectured that excessive SET binding to PP2A-C might pose some effect on the formation of PP2A-A/B/C holoenzyme.

**Figure 3.**
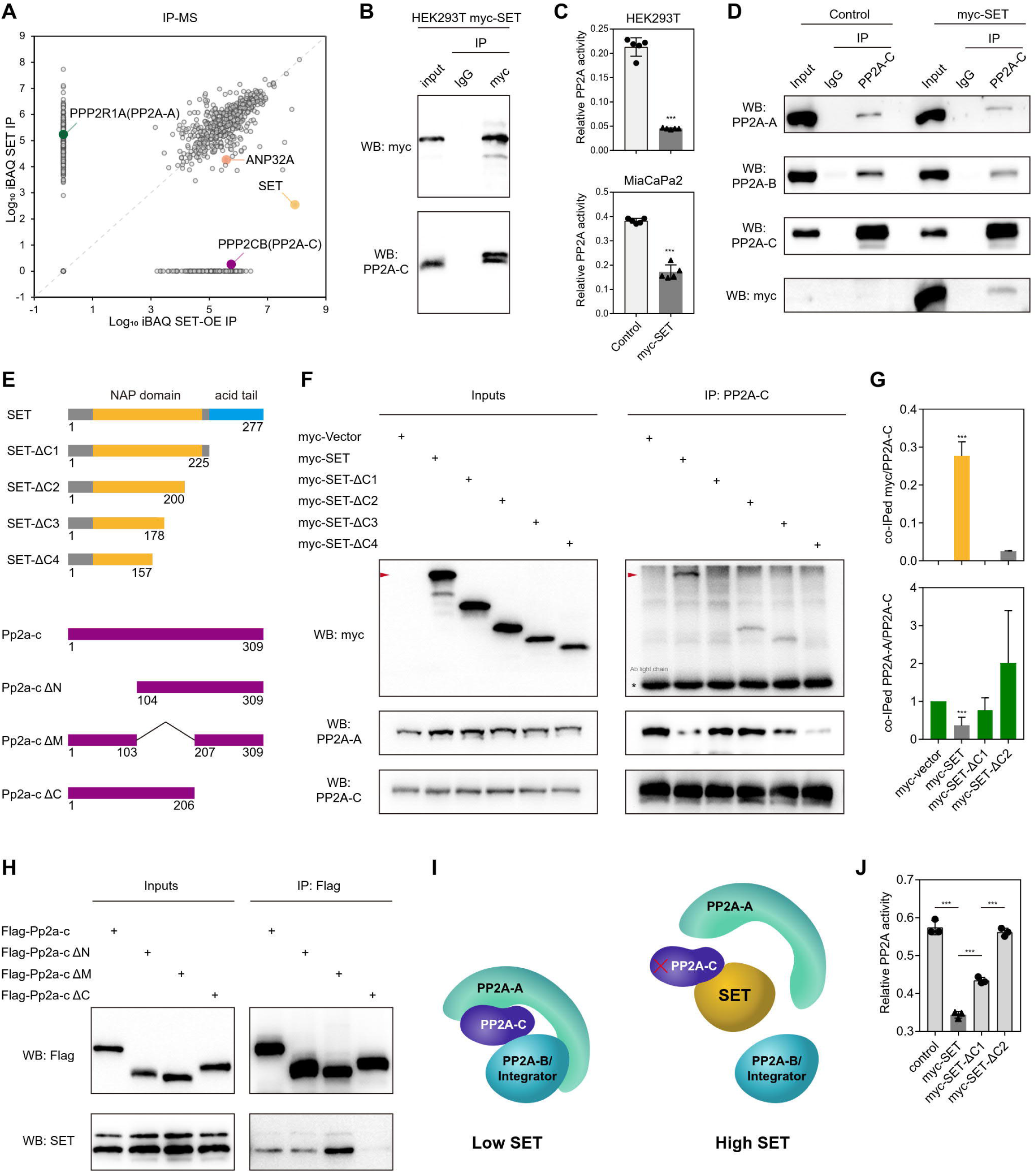
SET inhibits PP2A activity via disrupting PP2A-A/C complex. (A) Log_10_ iBAQ protein scores for SET IP mass spectrometry (MS) experiments in HEK293T SET-OE cells versus control HEK293T cells. ANP32A (PP32) is known to associate with SET in the INHAT complex. (B) Western blot of the co-IP product in the HEK293T SET-OE cells. (C) Gross PP2A activity was measured in the control or SET-OE cells. (D) Western blot of the co-IP product in the HEK293T control or SET-OE cells. (E) Illustration of the truncated mutations of SET and Pp2a-c protein. (F) Western blot of the co-IP product in the HEK293T cells stably transfected with wild-type SET or various truncated mutations. And the blotting bands were quantified using ImageJ software to calculate the relative amount of the co-IPed myc-SET as (G) in the upper panel, or the co-IPed PP2A-A by PP2A-C in the (G) lower panel. (H) Western blot of the co-IP product in the HEK293T cells stably transfected with wild-type Pp2a-c or various truncated mutations. (I) Mode of action of SET to inhibit PP2A activity. (J) Gross PP2A activity was measured in the HEK293T cells stably transfected with wild-type SET or different truncated mutations.

We constructed a series of truncated or deleted mutations in both SET and PP2A-C proteins (**Figure 3E**), trying to map the interaction domains in SET and PP2A-C. Full length SET bound to PP2A-C and interrupted the PP2A-A/C interaction (**Figure 3F**, arrow head). However, if a stretch of C-terminal domains in SET protein were deleted (SET-ΔC1 or SET-ΔC2), the truncated SET would hardly bind PP2A-C (**Figures 3F** and **3G**). Meanwhile, neither SET-ΔC1 nor SET-ΔC2 mutation would disrupt PP2A-A/C interaction (**Figures 3F** and **3G**). These results highlighted the importance of SET C-terminal domains in binding PP2A-C and impeding PP2A-A/C interaction. Using similar method, we mapped a PP2A-C region (aa207-309) that is critical for PP2A-C binding with SET (**Figure 3H**). Applying AlphaFold2-based protein complex simulations, we predicted a structural model of SET: PP2A-C interaction (**Figure S3D**). In this model, an evolutionarily conserved domain just adjacent to the SET C-terminal acid tail directly binds to a conserved region of PP2A-C (**Figures S3D** and **S3E**). According to this model, the SET bound with PP2A-C would induce a steric effect that interferes with PP2A-A/C interaction (**Figures S3D** and **3I**). PP2A-A/C interaction is the foundation for the complex of PP2A/B/C holoenzyme and full PP2A activity. So, it is possible that SET inhibits PP2A activity through disrupting the PP2A-A/C interaction. In accordance with this model, SET mutations with deletion of C-terminal domains showed substantially decreased binding with PP2A-C and no longer disrupted PP2A-A/C interaction or full PP2A activity (**Figures 3G** and **3J**). Our experiments suggested the importance of SET C-terminal domains in the stabilization of the binding with PP2A-C (**Figure S3F**).

### SET supports Pol II CTD phosphorylation through blocking PP2A activity

PP2A dephosphorylates many key factors that regulate transcription including the CTD domain of Pol II. If SET is a functional inhibitor of PP2A, it will maintain the phosphorylation of PP2A substrates. In the MS-based screen comparing the phospho-proteins in SET-KD versus the control cells (**Figure 4A**) (Kauko et al., 2020), we found that SET loss-of-function specifically decreased the phosphorylation of Pol II CTD domain in the fifth serine positions (Pol II S5-p), though western blot suggested that SET-KO would decrease Pol II S2-p and S7-p as well (**Figure 4B**). Pol II S2, S5 and S7 are phosphorylated by CDK7 or CDK9 and the fully phosphorylated Pol II is activated in transcription (Allen and Taatjes, 2015; Jonkers and Lis, 2015; Peterlin and Price, 2006; Zhou et al., 2012). CDK9 inhibition largely down-regulated the phosphorylation of these serine and SET-KO showed a similar function (**Figure S4A**). CDK9 inhibition plus SET-KO, however, showed no additive effect to decrease the Pol II CTD phosphorylation (**Figure S4A**), which implies that SET and CDK9 work in the same pathway.

**Figure 4.**
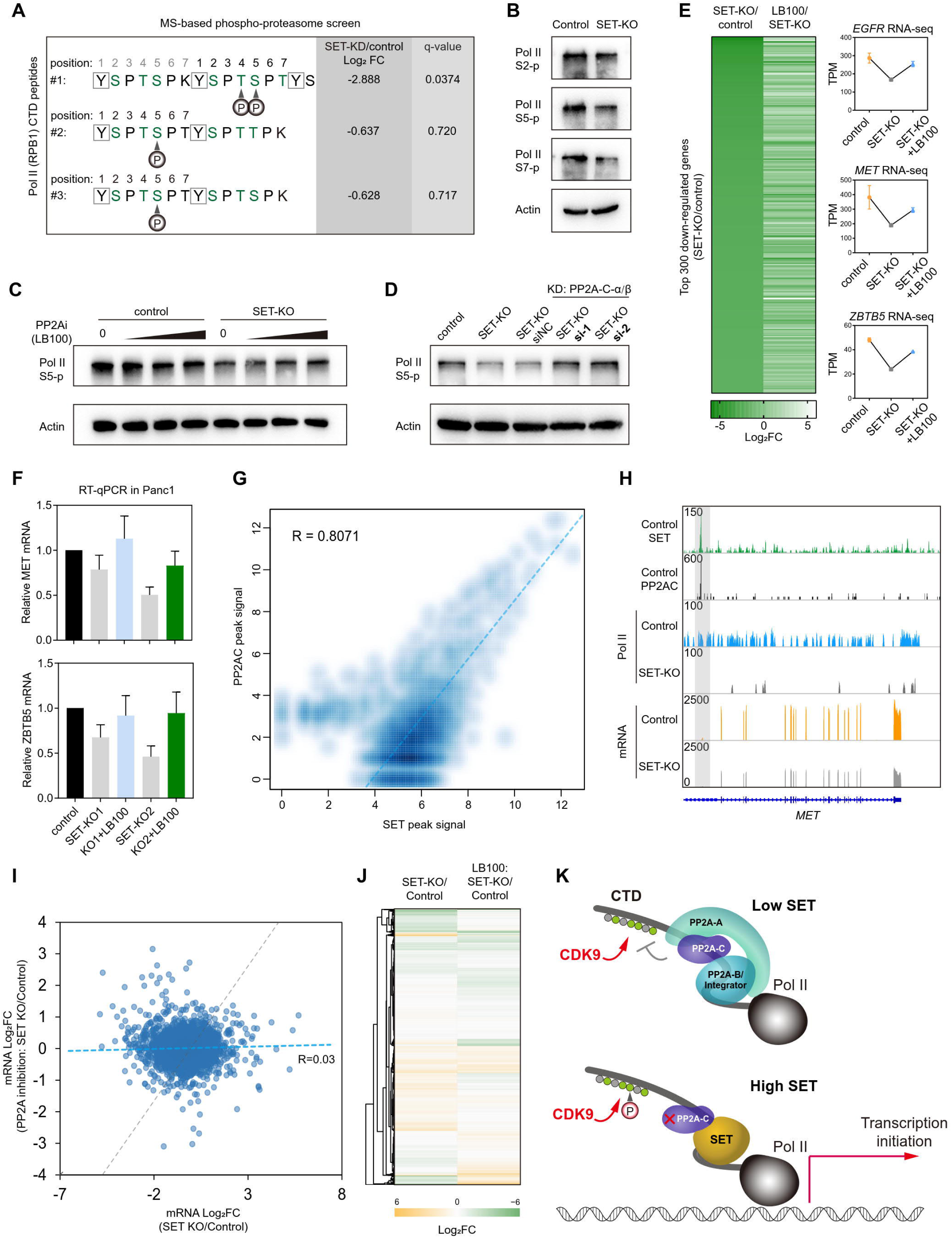
SET supports Pol II CTD phosphorylation through blocking PP2A activity. (A) TiO_2_ enrichment and mass spectrometry assays indicates the phosphorylated sites in Pol II CTD and their fold change after SET knock-down. (B) Western blot of the cell extract of BXPC3 control and SET-KO cells. (C) Western blot of the cell extract of BXPC3 control and SET-KO cells upon treatment with PP2A inhibitor. (D) Western blot of the lysates of different cells under indicated treatments.si-1/2 indicates two cocktails of siRNAs targeting both PP2A-C - α and- β . (E) left panel: Heatmap shows the top 300 down-regulated genes in BXPC3 SET-KO cells v.s. control cells and their fold change after PP2A inhibition by LB100; right panel: Representative genes and their mRNA levels in different situations. (F) Representative genes and their mRNA levels quantified via RT-qPCR in different situations. (G) Density plot shows the relationship between SET and PP2A-C peak signal along the genome. (H) Tracks profile the SET and PP2A-C chromatin distribution in BXPC3 cells, as well as Pol II occupancy/mRNA level changes after SET knock-out in a representative locus. (I) Scatter plot and (J) Heatmap shows the fold change of DEGs in SET-KO cells v.s. control cells with or without inhibition of PP2A activity. (K) Model diagram explains how SET supports the CDK9-induced, Pol II CTD phosphorylation dependent activation of gene transcription, through directly inhibition of PP2A.

Wondering whether SET maintains Pol II CTD phosphorylation via blocking the PP2A activity, we observed Pol II S5-p levels in the control and SET-KO cells with or without PP2A inhibition. In the control cells, where SET level was high and PP2A was expected to be already blocked, PP2A inhibitor LB100 could hardly increase Pol II S5-p any further; while in the SET-KO cells where PP2A activity was high and Pol II S5-p was low, PP2A inhibition increased the level of Pol II S5-p (**Figure 4C**). We knocked-down both PP2A-Cα and β isoforms (**Figure S4B**) in the SET-KO cells and also observed the elevation of Pol II S5-p level (**Figure 4D**). These results demonstrated that SET maintains Pol II S5-p level via blocking PP2A activity.

To investigate whether SET drives the massive hyper-transcription through disrupting PP2A, we analyzed the effects of PP2A inhibition on mRNA transcriptome in both control and SET-KO cells. We found that, for the most decreased oncogenes in the SET-KO cells including *EGFR*, *MET* and *ZBTB5*, PP2A inhibition will increase their mRNA levels (**Figures 4E** and **4F**). In fact, for all the down-regulated genes in SET-KO cells versus the control cells, 67.5% of them were eventually up-regulated by PP2A inhibition (**Figure S4C**). It means that PP2A might play a great part in the SET induced transcriptome remodeling. PP2A-C CUT&Tag data showed that PP2A-C not only bound with SET but also co-localized with SET along the genome (**Figure 4G**). PP2A-C and SET even co-targeted the similar genome DNA motifs (**Figure S4D**).

These facts highlighted the possibility that SET could support Pol II function and mRNA transcription by in situ inhibition of chromatin associated PP2A (**Figure 4H**). Moreover, in the LB100 treated cells where PP2A activity was blocked, SET-KO could no longer induce the transcriptome phenotype as it did in the PP2A-normal cells (**Figures 4I** and **4J**). In conclusion, SET prompts global transcription at least partially by disrupting PP2A holoenzyme and PP2A activity (**Figure 4K**).

### SET suppresses promoter histone acetylation, while maintaining the genome-wide acetyl-histone landscape and enhancer activation in an “offshore” manner

SET was identified as a member of the INHAT (inhibitor of histone acetyltransferase) complex (Seo et al., 2001) and the insulator of histone acetylation (Wang et al., 2016). Deeper exploration of the SET interactome data also found CTBP2 and HDAC1, two key components of histone deacetylase complexes, as possible binding partners of SET (**Figures 5A** and **5B**). Considering that histones are also stable binding targets of SET (**Figure S5A**), SET is very likely to pose an active impact on histone acetylation. In fact, we have validated that SET-KO increased the overall acetylation level of histone H3 and H4, while SET-OE suppressed the histone acetylation (**Figure 5C**).

**Figure 5.**
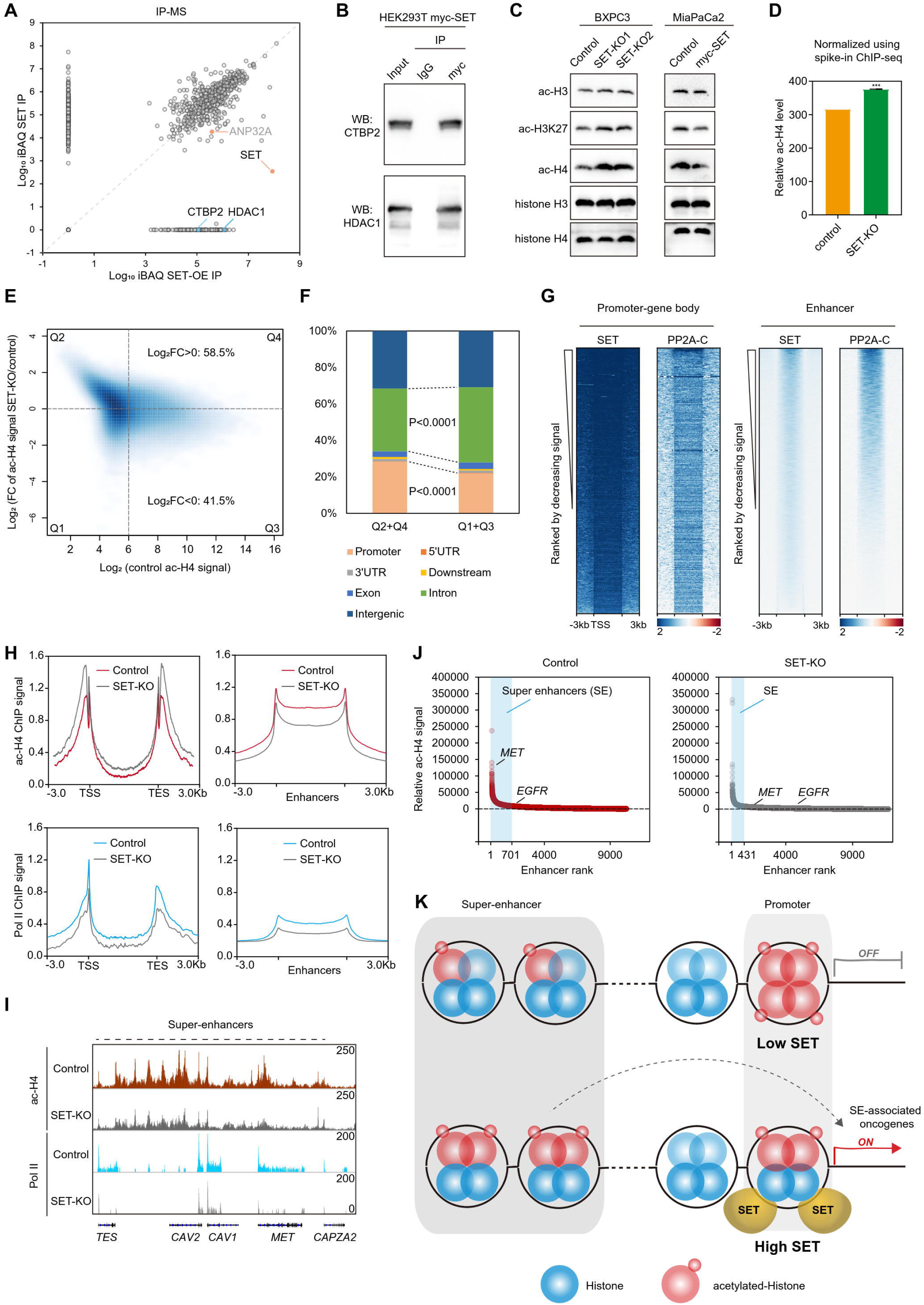
SET suppresses promoter histone acetylation, maintains the genome-wide acetyl-histone landscape and enhancer activity. (A) Log_10_ iBAQ protein scores for SET IP mass spectrometry (MS) experiments in HEK293T SET-OE cells versus control HEK293T cells, the same data as in Figure 3A but different data spots are labeled. (B) Western blot of the co-IP product in the HEK293T SET-OE cells. (C) Western blot of the histone extract of the indicated cells. (D) The same ratio of spike-in drosophila chromatin was added to the BXPC3 cell chromatin before ChIP, the relative ac-H4 level was calculated using the formular: Relative level= (Human genome specific reads) / (Drosophila genome specific reads) -1. (E) Density plot showing the log_2_ fold change of ac-H4 signal after SET-KO v.s. signal in the control sample. Regions are divided into four classes on the basis of fold change and signal in DMSO treatment. (F) Association of regions from (E) with genomic features compared with an overall genomic distribution. Note promoters (−3 kb from TSS) and downstream (+3 kb from TTS). (G) Heatmaps show the SET and PP2A-C CUT&Tag signals at scaled gene or enhancer regions ranked by decreasing occupancy. (H) Metaplots show the ac-H4 and Pol II ChIP-seq signals at scaled gene or enhancer regions in the control or SET-KO cells. (I) Tracks profile the ac-H4 intensity and Pol II occupancy at a representative super-enhancer region in the control or SET-KO cells. (J) Scatterplot depicts distribution of ac-H4 ChIP-seq density and ranking across all enhancers in the control or SET-KO cells. Super-enhancers (SEs) were identified with ROSE program. Enhancers (including SEs) were associated with the nearest gene within 50 kb. (K) The model diagram depicts that SET prefers to bind gene promoters while allowing enhancer to accumulate histone acetylation and to activate enhancer-associated oncogene transcription.

SET-KO induced hyper-acetylation of histone H4 could be verified using quantitative ChIP-seq (**Figure 5D**). These findings brought the doubt why a histone acetylation insulator could reversely stimulate the global hyper-transcription? Although we have provided evidence that SET could induce massive hyper-transcription via inhibiting PP2A, it is still necessary to study the detailed impacts of SET on genome-wide histone acetylation.

Although bound with both histone H3 and H4 (**Figure S5A**), SET showed stronger impact on histone H4 acetylation level (**Figure 5C**). It has also been suggested that histone H4 acetylation, which is highly correlated with histone H3K27 acetylation, is another marker of enhancers (Slaughter et al., 2021). So, we chose to study the genome-wide changes in histone H4 acetylation after SET loss-of-function. After systematic detection of histone H4 acetylation landscapes in the control and SET-KO cells (**Figure 5E**), we found the histone H4 acetylation landscape was “depolarized” after SET-KO, which means those genome loci with a lower level of histone H4 acetylation tended to increase acetylation after SET-KO, while the loci with a high level of acetylation were to decrease (**Figure 5E**, Q1 and Q2). On the whole, more loci (58.5%) increased their acetylation level after SET-KO. But the changes of histone acetylation in distinct genome elements are different. Generally, the promoter regions were inclined to gain histone H4 acetylation but the introns tended to lose H4 acetylation after SET-KO (**Figure 5F**).

These findings prompt us to study the genome element-specific binding pattern of SET. We noticed an obvious phenomenon that SET preferred to bind gene promoters rather than enhancers (**Figure 5G**). In accordance with this binding pattern, SET-KO induced histone hyper-acetylation was mainly concentrated in the promoter-gene body but not in the enhancer regions (**Figure 5H**). We also found in the SET-KO cells that histone H4 hyper-acetylation at gene promoters was not related to the up-regulation of gene expression (**Figure S5C**). On the contrary, the down-regulated genes after SET-KO could have elevated histone H4 acetylation in their promoter or gene body, compared with the up-regulated genes (**Figures S5C** and **S5D**).

The SET-KO cells exhibited down-regulated histone H4 acetylation with decreased Pol II occupancy in the enhancers and super-enhancers (**Figures 5H**, **5I** and **S5E**), and less enhancer RNA (eRNA) transcribed from the super-enhancers (**Figure S5E**). All these indicated the inactivation of enhancers. Compared with the control cells, SET-KO cells had fewer number of super-enhancers (**Figure 5J**). Many super-enhancers in the control cells are associated with RTK or AKT signaling pathway genes (including *MET* and *EGFR*, **Figure S5F**). When the super-enhancer activity was attenuated by SET-KO, those RTK genes with other growth-promoting genes were also turned down (**Figures 5J, S5G and S6**). For the genes related to the RTK, AKT pathway or positively regulating cell proliferation (shown in **Figure S5G**) which were down-regulated after SET-KO, we analyzed their related enhancer regions and observed that most sub-enhancer loci with decreased ac-histone H4 after SET-KO actually lacked SET binding in the control cells (**Figure S6**). All the phenomena above support the model that higher level of SET prefers binding to the histones of gene promoters rather than the enhancers, so the excessive SET insulated the gene promoters from further acetylation while allowing the super-enhancer regions lacking SET binding to accumulate acetylation markers and be activated (**Figure 5K**).

Compared with gene promoters, the super-enhancers play a more prominent role in orchestrating oncogene expression selectively (Loven et al., 2013). So, in the cells with higher level of SET, the super-enhancer histone acetylation evaded from SET insulation seemed to overweigh the promoter hypo-acetylation by SET, to ultimately induce massive hyper-transcription of growth-essential oncogenes. Although not regulating enhancer directly, SET seems to permit super-enhancer activation in an “offshore” manner.

### High level of SET is correlated with Pol II CTD hyper-phosphorylation, c-MET over-expression and hyper-sensitivity to the c-Met inhibitor in cancers

Loss of SET attenuated the hyper-transcription of growth-essential genes and suppressed tumor growth. But SET in itself is not a druggable target in cancers. In search of the Achilles’ heel in SET over-expressing cancers, we analyzed the correlation between drug sensitivity and SET expression level in cancer cell lines (**Figure 6A**). Statistically, cancer cells with elevated SET showed increased vulnerability to compounds with various cellular targets (**Figure S7A**), especially the Taxels and c-Met inhibitor Tivantinib (**Figures 6A**, **6B** and **S7A**). RTK c-Met is encoded by oncogene *MET*, which locates in a super-enhancer region substantially suppressed by SET-KO (**Figure 5I**). It’s noteworthy that SET and PP2A-C also co-targeted the promoter of a *MET* variant which encoded a short isoform of c-Met (**Figure 4H**). In concert, MET mRNA level was significantly suppressed by SET-KO and partially restored by PP2A inhibition (**Figure 4E**). We further validated that c-Met protein level was decreased by SET-KO (**Figure 6C**). Comparing to the SET-KO cells, the control cells were more sensitive to the c-Met inhibitor (**Figures 6D** and **6E**). These observations demonstrated that MET is prominent among the massive hyper-transcribed genes driven by SET.

**Figure 6.**
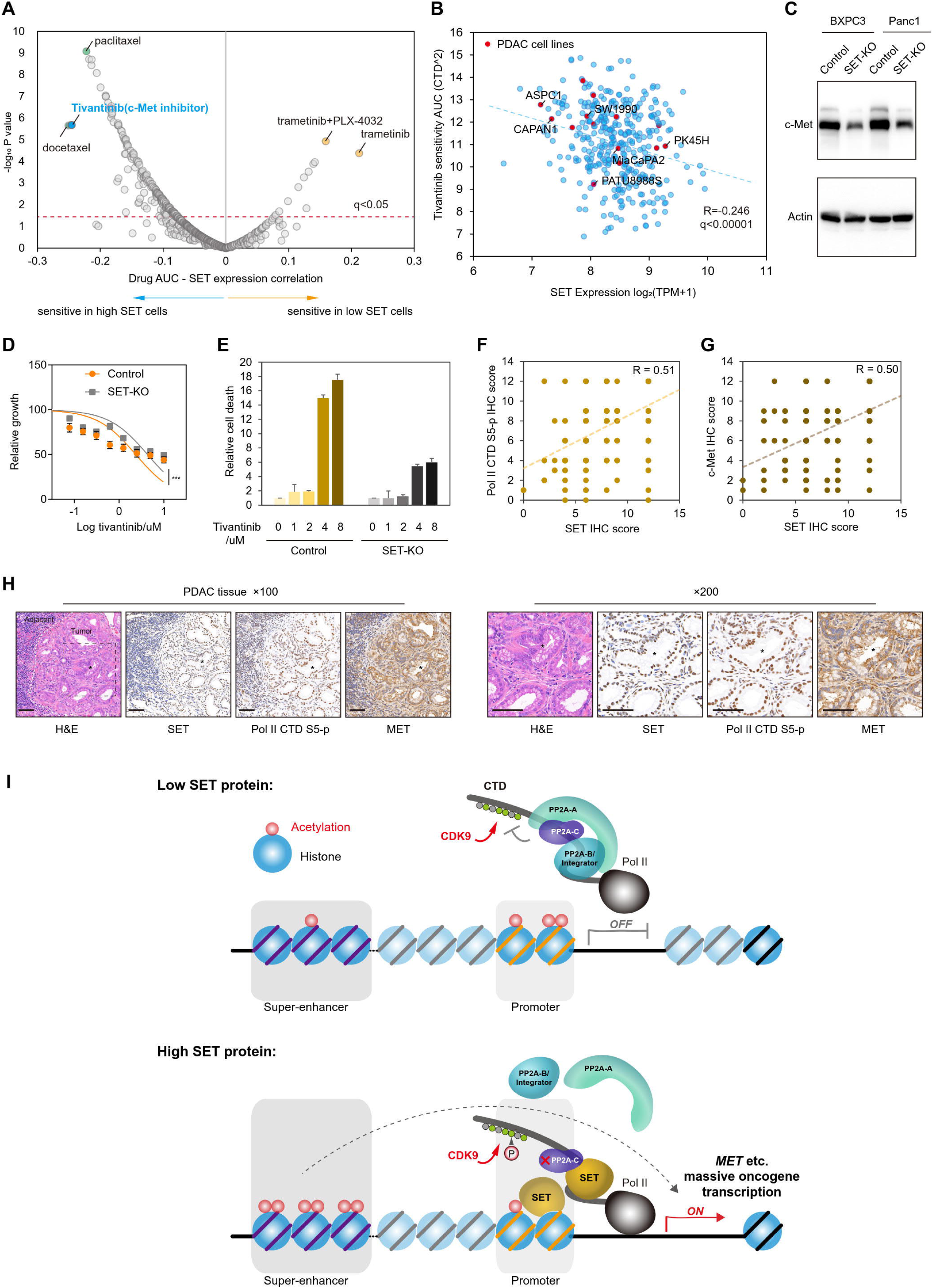
High level of SET is correlated with Pol II CTD hyper-phosphorylation, c-MET over-expression and hyper-sensitivity to the c-Met inhibitor in cancers. (A) Scatter plot shows the relationship between SET expression and drug sensitivity in the CCLE cancer cell lines. (B) Scatter plot shows the relationship between SET expression and cell sensitivity to the c-Met inhibitor tivantinib in multiple cancer cell lines. (C) Western blot of the cell extract of the control and SET-KO cells. (D) Drug IC_50_ assay was carried out in the control and SET-KO Panc1 cells. (E) Cell death assay after c-Met inhibitor treatment was carried out in the control and SET-KO BXPC3 cells. (F) Scatter plot indicates the correlation between SET IHC score and Pol II CTD S5p IHC score or (G) c-Met IHC score in PDAC tissues. (H) Representative IHC images, scale bar: 50μ The star marks the same position in the tissue. (I) The model diagram of this study.

Pol II CTD S5-p is a hallmark of Pol II activation and transcription initiation (Chipumuro et al., 2014; Thandapani, 2019), our experiments have suggested that SET supported Pol II S5-p and the hyper-transcription in PDAC cells. In our cohort of PDAC patients, we found Pol II S5-p level was increased in the PDAC tissues (**Figure S7B**). High Pol II S5-p level correlated with shorter survival time in PDAC (**Figure S7H**), reflecting a massive hyper-transcription program favoring the expression of growth-essential genes.

Meanwhile, *MET* expression was also elevated in PDAC (**Figure S7C**) and the increased *MET* (c-Met) is positively correlated with Pol II S5-p level and SET level (**Figures 6F**, **6G**, **S7D-S7G**), implying that up-regulation of oncogene *MET* is part of the hyper-transcription program prompted by SET. Moreover, we observed that *MET* high expression is correlated with short survival time of PDAC patients, indicating that *MET* over-expression contributed to the malignancy of PDAC (**Figures S7I and S7J**) in accordance with previous reports (Ebert et al., 1994; Kiehne et al., 1997; Lee et al., 2003; Qin et al., 2022; Yan et al., 2018). In conclusion, IHC study in the PDAC consecutive sections (**Figure 6H**) supported the hypothesis that over-expressed SET, supporting CDK9-induced Pol II CTD phosphorylation, prompts the hyper-transcription of growth-essential genes including the oncogene *MET* (**Figure 6I**).

## Discussion

In this study, we have observed a synergetic co-operation of SET protein’s multiple functions in regulating global transcription. As a disruptor of PP2A activity, SET is able to prompt CDK9-induced and Pol II-mediated transcription, in site; and as a promoter-philic histone acetylation insulator, SET selectively allows the transcription of super-enhancer related oncogenes, offshore. In a concerted way, these two distinct functions of SET work together without confliction (**Figure 6I**).

Our results indicated that SET disrupted PP2A-A/C interaction and abolished the dephosphorylation of Pol II CTD, especially on serine 5, by PP2A. So, it is explainable that SET supported CDK9 to phosphorylate Pol II CTD hence to activate mRNA transcription. It has been demonstrated that CDK9 and PP2A antagonize each other to regulate the phosphorylation on Pol II CTD (Tellier et al., 2022; Vervoort et al., 2021; Zheng et al., 2020). Pol II S5-p was reported to be related to transcription initiation while S2-p is more significant in the transcription elongation after Pol II pausing (Eick and Geyer, 2013; Larochelle et al., 2012; Zhou et al., 2012). Though our results showed that SET-KO induced both significant decrease of Pol II S5-p and moderate down-regulation of S2-p, we didn’t observe obvious differences in Pol II pausing in SET-KO cells versus the control cells. So, it seems that SET loss-of function is not 100% identical with PP2A gain-of-function. We speculate that SET-loss suppressed transcription initiation so profoundly, which overwhelms its effects on transcription elongation and other down-stream events. And it also implies that PP2A inhibition is only part of the mechanism underlying SET-induced hyper-transcription. Actually, in SET-KO cells both the phosphorylated and non-phosphorylated Pol II were decreased compared to the control cells (**Figure S4A**), which means SET positively regulates both the phosphorylation level and the total protein level of Pol II. But we are not clear how SET regulates Pol II (Rpb1) protein level because we have not found direct interaction between SET and Rpb1. Moreover, SET was reported to be a histone chaperone with possible chromatin remodeling activity (Gamble et al., 2005; Gamble and Fisher, 2007; Muto et al., 2007; Okuwaki and Nagata, 1998). We suspect that SET-KO might also induce chromatin restriction to directly hinder the Pol II engagement to the chromatin. This effect could overlap with SET’s function as a PP2A inhibitor, to jointly impact the transcription.

SET has been identified as an endogenous PP2A inhibitor, but the mechanism of SET inhibiting PP2A is not fully elucidated (Arnaud et al., 2011; Saito et al., 1999). Through quantitative IP-MS, competitive binding assay and sub-domain mapping, we suggest that SET inhibits PP2A by directly binding to PP2A-C and disrupting PP2A-A/C interaction. Our experiments indicated that SET C-terminal acid tail is indispensable for its binding with PP2A-C and the ability to disrupt PP2A-A/C, while the predicted structural model showed that SET-PP2A-C interaction is mediated by a conserved domain just up-stream adjacent to the acid tail of SET. The structural model admitted that SET acid tail owns a flexible structure difficult to certify the precise conformation, which we speculate led to the discrepancy between the theoretical model and experimental results. The acid tail of SET might be prerequisite for the direct binding sites in SET and PP2A-C to recognize each other.

Especially, we found that oncogene *MET* is one of the important down-stream targets of SET induced hyper-transcription. Relative study in the PDAC patient cohort and previous studies all suggested that *MET* might contribute to the malignancy of PDAC (Kitajima et al., 2008; Lux et al., 2019; Noguchi et al., 2018; Yan et al., 2018). Our results added to the pharmaceutical value of c-Met inhibitors in the treatment of PDAC (Herreros-Villanueva et al., 2012; Jin et al., 2008; Jin et al., 2021; Li et al., 2011; Mori et al., 2021).

We observed that SET preferred binding to gene promoters rather than enhancers, which explained why SET, as a histone acetylation insulator, allowed the enhancers to accumulate histone acetylation. But what navigated SET to the gene promoters specifically? Considering that PP2A complex was found to engage chromatin at gene promoters (Zheng et al., 2020), and that PP2A-C not only bound SET but also co-localized with SET along the genome, we speculate that SET might be tethered to gene promoters by chromatin-associated PP2A. However, we couldn’t rule out other factors that determine the genome target specificity of SET and the question how SET evades the enhancers remains open.

### Limitations of the study

In the SET-KO cells, we found Pol II (Rpb1) total protein level and CTD phosphorylation level were both decreased compared with the control cells. We explained that SET maintains Pol II CTD phosphorylation level via inhibiting PP2A but we didn’t study the mechanism that SET maintains Pol II total protein level. We found that SET is important for Pol II function and transcription initiation. But we could not certify the effects of SET on transcription elongation and other down-stream events, because SET-KO induced a profound loss of Pol II engagement to the chromatin that might mask other down-stream events in RNA transcription. We found that SET maintains enhancer activation and we have known that SET prefers binding to gene promoters rather than enhancers which allows the enhancer regions to gain histone acetylation, but we didn’t investigate whether SET stimulates enhancers actively or directly. Moreover, it is ideal to re-activate PP2A in SET-OE cells hence to further verify whether SET induced hyper-transcription or cell growth was mediated by PP2A inhibition, however these results cannot be achieved because phenothiazine based PP2A activating compounds iHAP1 and DT-061 had recently been found to possess PP2A-independent cytotoxicity (Vit et al., 2022).

## ACKNOWLEDGMENTS

We thank Professor Xudong Wu of Westlake University for the SET-PP2AC complex structure simulation, and Professor Fei Xavier Chen of Fudan University Shanghai Cancer Center for the sharing of nascent RNA-seq technology. This work was funded by National Natural Science Foundation of China (U21A20374, 82072698, 82002541, 82203434), China Post-doc science Foundation (2021M690698), Shanghai Municipal Science and Technology Major Project (21JC1401500), Scientific Innovation Project of Shanghai Education Committee (2019-01-07-00-07-E00057), Clinical Research Plan of Shanghai Hospital Development Center (SHDC2020CR1006A), Xuhui District Artificial Intelligence Medical Hospital Cooperation Project (2021-011), Shanghai Rising-Star Program (no. 20QA1402100)

## Author contributions

D.W. and S.S conceptualized the study. D.W. and H.X. found the connectivity between SET over-expression and CDK9 gain-of-function, as well as the positive effect of SET upon Pol II-mediated transcription and super-enhancer activation. J.X. provided experiment platform and patient samples. H.X. performed patient studies, cell culture and animal experiments. D.W. H.X. and YB.L. performed the nascent RNA-seq, RNA-seq, ChIP-seq and CUT&Tag. D.W. performed the computational analysis. YL.L. drawn the heatmaps of RNA-seq. D.W. wrote the manuscript. D.W., X.Y. and S.S. supervised the work.

## Figure legends

## STAR METHODS

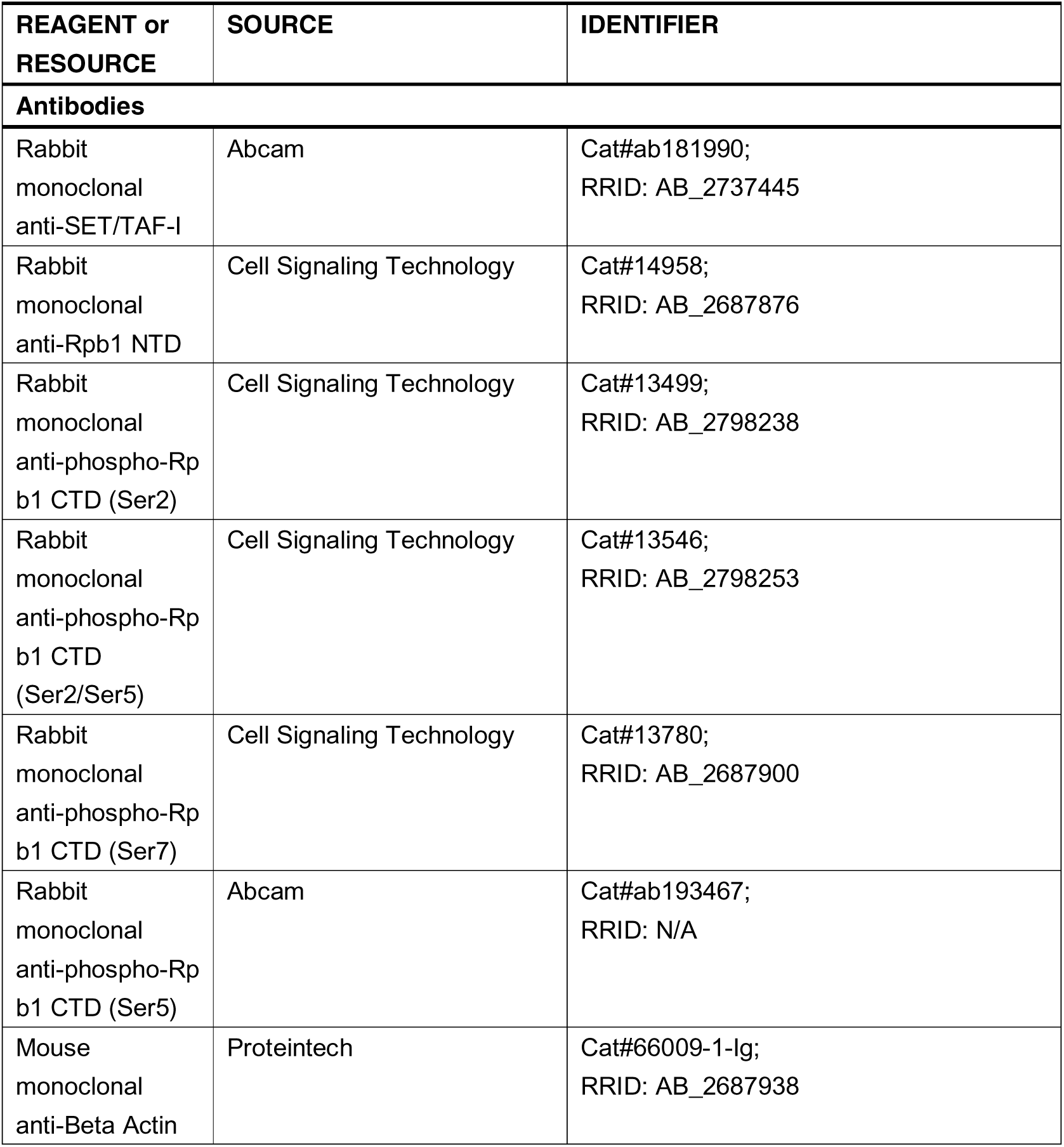
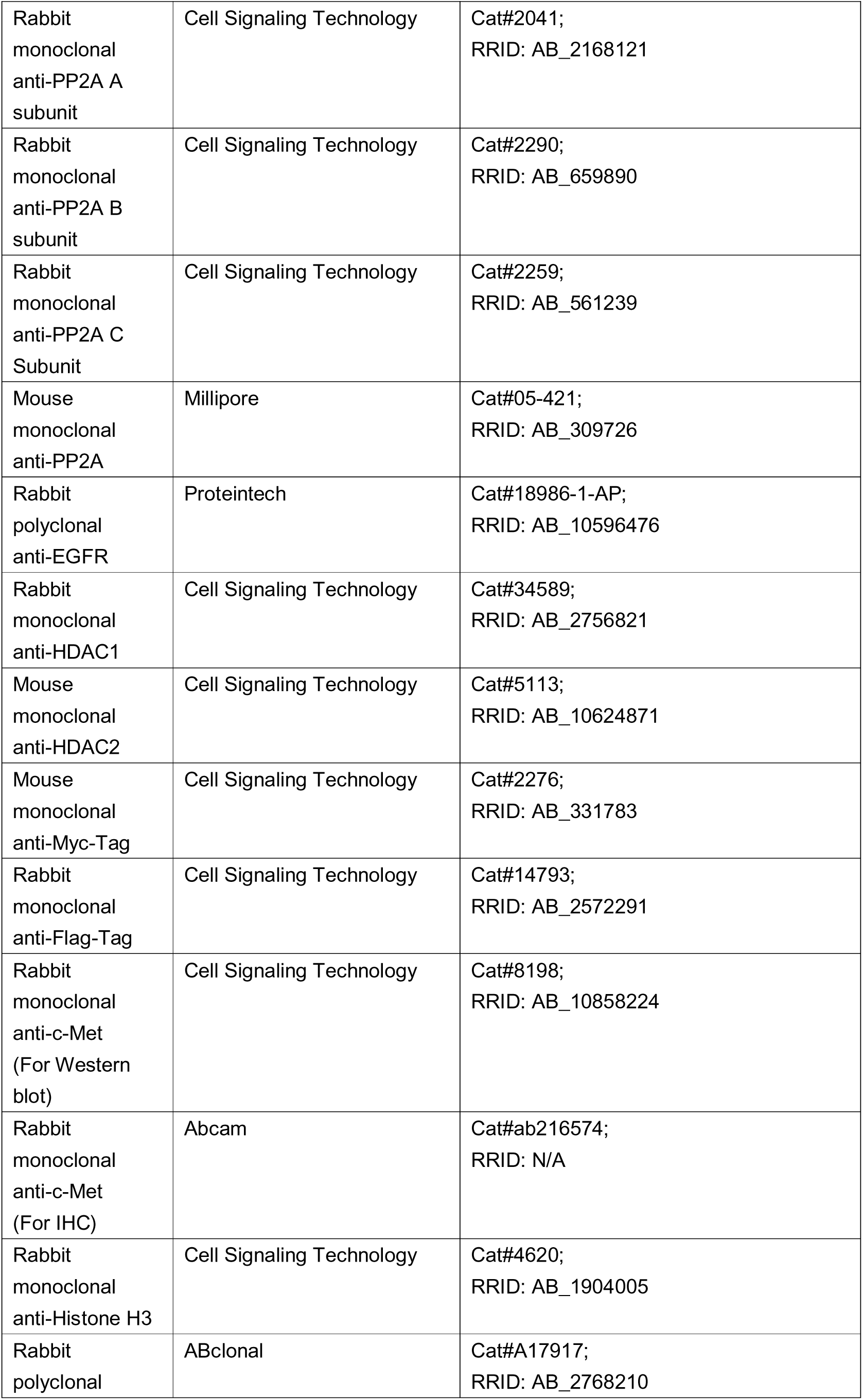
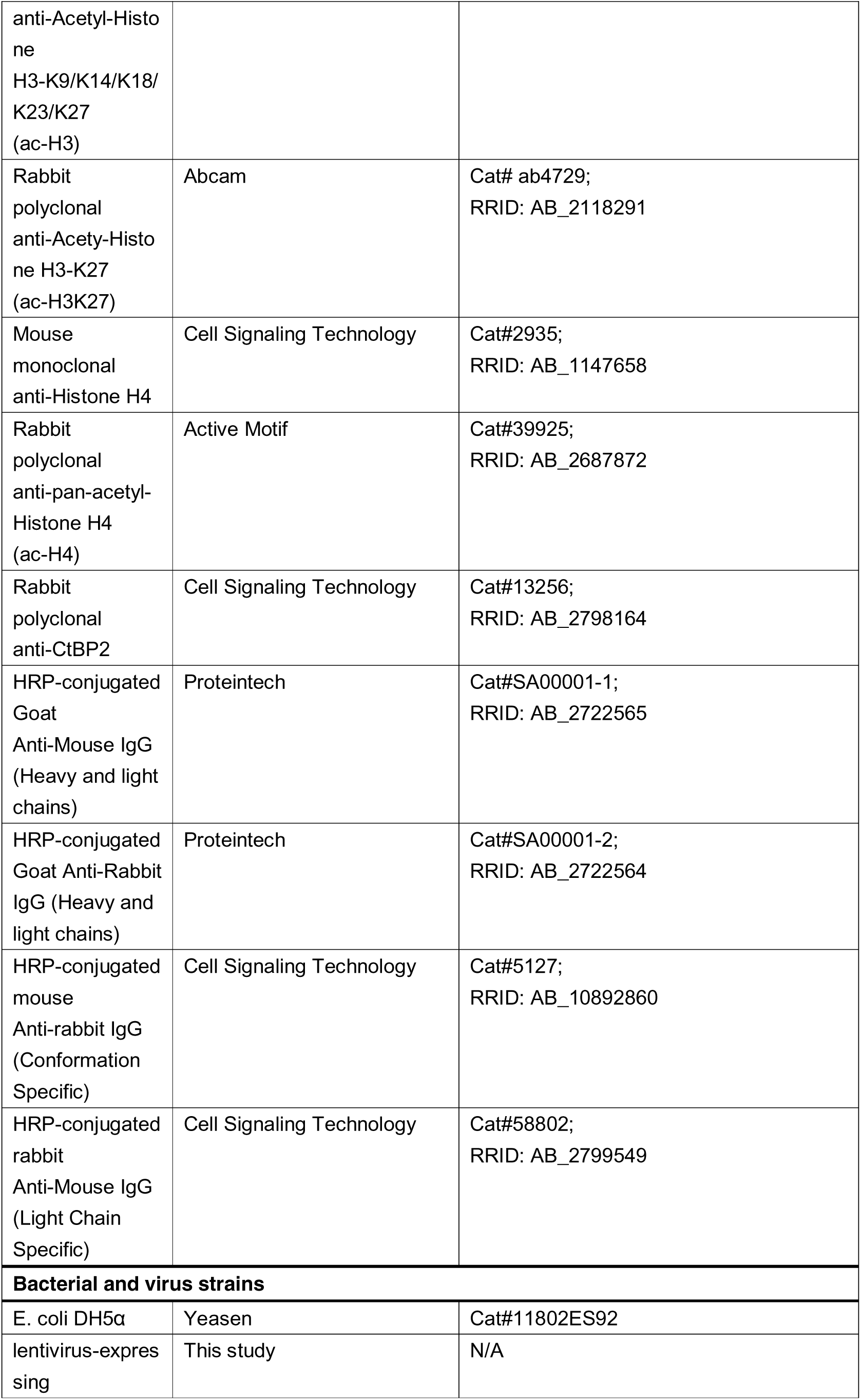
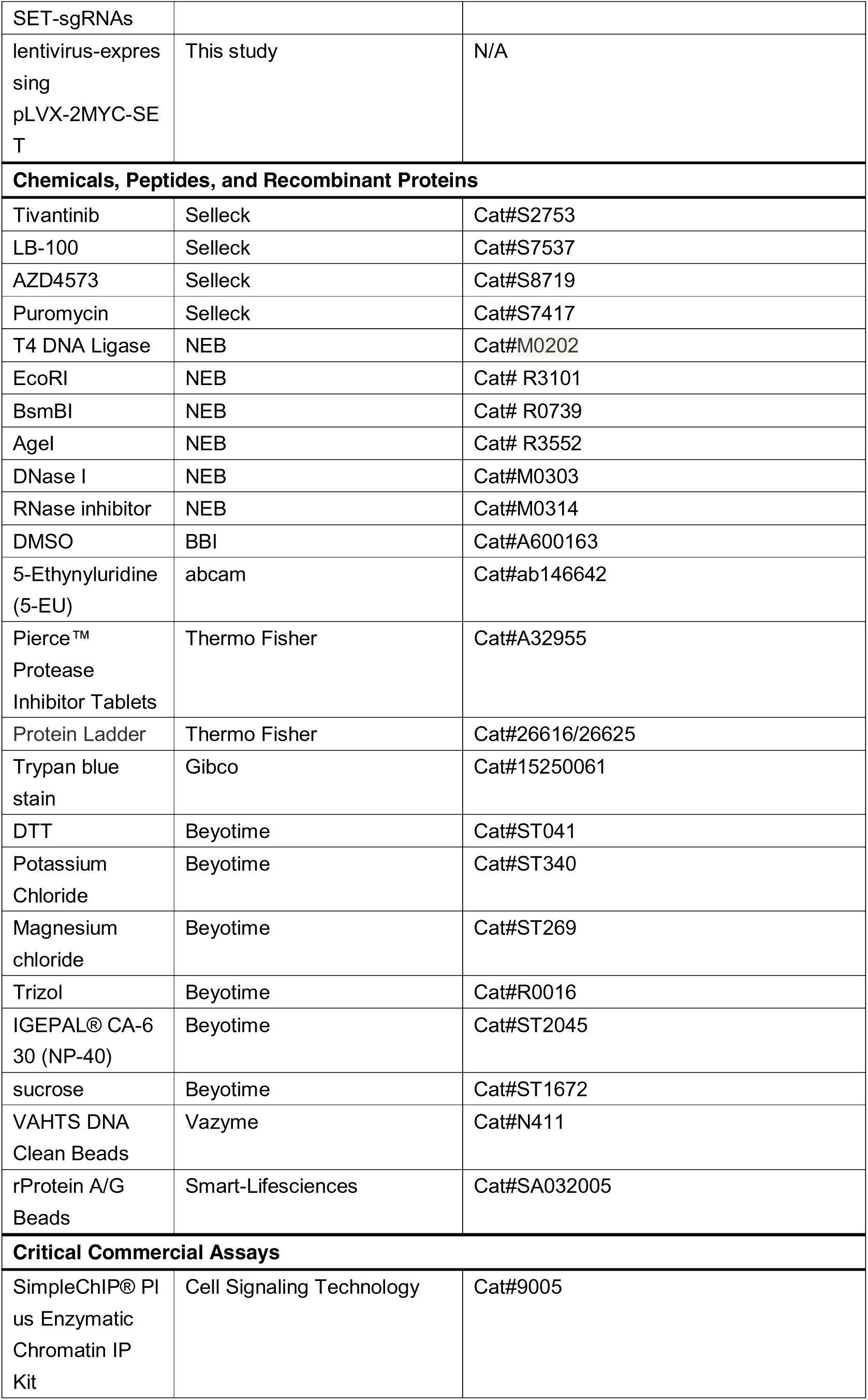
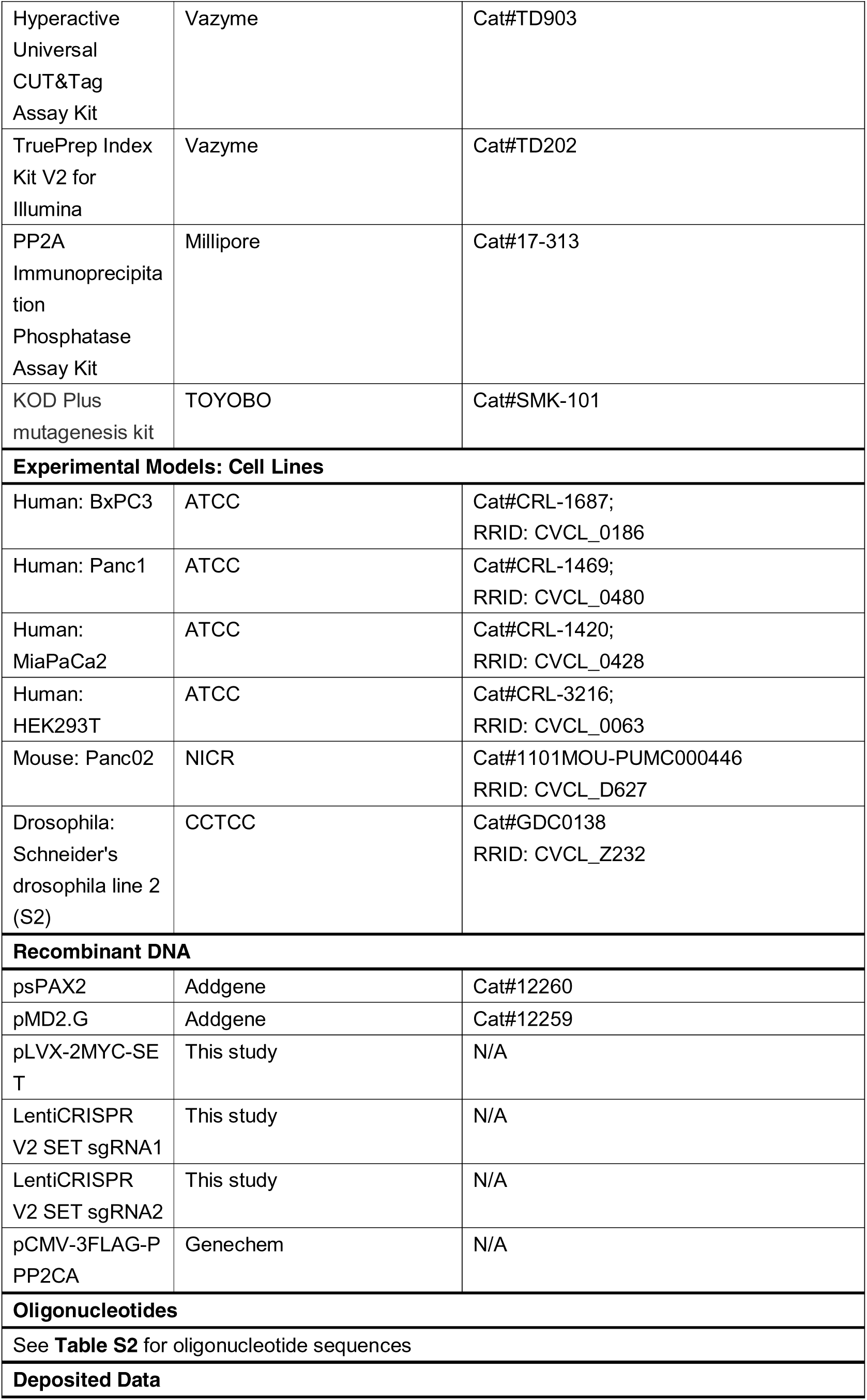
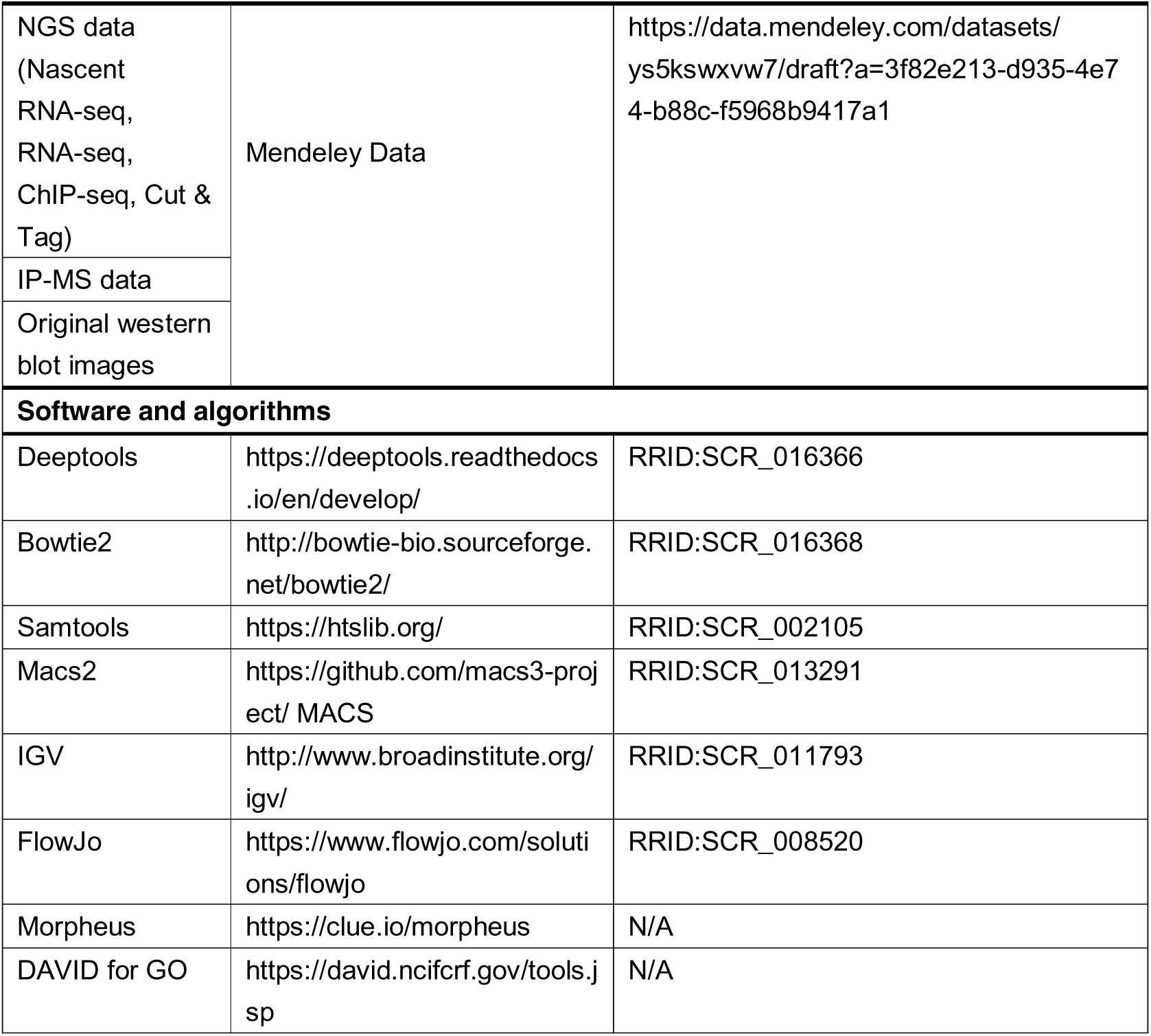
KEY RESOURCES TABLE

## RESOURCE AVAILABILITY

### Lead Contact

Further information and requests for resources and reagents should be directed to Si Shi

### Materials Availability

Cell lines and plasmids are listed in the key resources table. Oligonucleotides are listed in **Table S2**.

### Data and Code Availability

All sequencing tracks generated from nascent RNA-seq, RNA-seq, ChIP-seq and CUT&Tag reported in this paper and gene expression TPM lists have been deposited to Mendeley Data (https://data.mendeley.com/datasets/ys5kswxvw7/draft?a=3f82e213-d935-4e74-b88c-f5968b9417a1) along with the IP-MS data and the original Western blot images. Any additional information required to reanalyze the data reported in this paper is available from the lead contact upon request.

## EXPERIMENTAL MODEL AND SUBJECT DETAILS

### Patient study

The paraffin section specimens were obtained from patients diagnosed with PDAC who underwent surgery at the Fudan University Shanghai Cancer Center (FUSCC) from 2016 to 2018. Totally the 239 patients, 104 had solely PDAC tissue, while the other 135 had both cancerous and adjacent tissue. PDAC RNA was extracted from the surgical tissues of 96 patients diagnosed in 2013 and 2014. Each patient provided an informed consent, and all experiments were conducted with the approval of the FUSCC Clinical Research Ethics Committee. The disease diagnosis and immunohistochemistry staining grading were performed by two experienced pathologists. The latest follow-up date was November 2020.

### Cell lines and culture conditions

We cultured and passages HEK293T, Drosophila S2, mouse pancreatic cancer cell line Panc02 and human pancreatic cancer cell lines BxPC3, PANC1, and MiaPaca2. The cells were identified by DNA fingerprinting. BxPC3 cells were cultured in RPMI 1640 supplemented with 10% FBS. Drosophila S2 cells were maintained with Schneider’s Drosophila medium with 10% FBS. The other cells were grown in DMEM with 10% FBS. All the media were supplemented with 100 U/mL penicillin and streptomycin. Cell lines were incubated at 37°C under 5% CO2 and a PCR test for mycoplasma is performed every two months.

### Mouse experiments

All animal experiments were conducted at the Animal Center of Fudan University Shanghai Cancer Center, and the Animal Ethics Committee approved the research. A barrier system with ventilated cages and automatic water dispensers was utilized to house mice. The animal house is programmed with a temperature range of 20–26°C, a humidity range of 40–60%, and a 12-hour day/night cycle. Female BALB/c nude mice (5–6 weeks of age, 18–22g, Gempharmatech) were injected subcutaneously with 1 × 10^6^ BxPC3 control or SET knockout cells. 1 × 10^6^ MiaPaCa2 control or SET over-expression cells were also injected in another group. The mice were sacrificed and the tumors were weighed 4 weeks after injection.

## METHOD DETAILS

### Immunohistochemistry (IHC)

The IHC analysis was conducted in the same manner as previously described (Hua et al., 2022). Briefly, tissue sections with a thickness of 3 um were deparaffinized in xylene and rehydrated in gradient alcohol solutions. Heat-mediated antigen retrieval was performed using EDTA buffer (pH 9.0) in a pressure cooker for 10 minutes. The slides were then cooled to room temperature and incubated with a 3% H_2_O_2_ solution for 20 minutes to inhibit endogenous peroxidase activity. After 2 hours of blocking with 3% normal goat serum, the slides were treated with optimum dilutions of primary antibodies at 4°C overnight. The samples were then stained using a two-step polymer-horseradish peroxidase method using DAB as the chromogen and Mayer’s hematoxylin as the counterstain. Scores for IHC staining were calculated by multiplying the proportion or area (0, <5%; 1, 5–25%; 2, 25–50%; 3, 50–75%; and 4, >75%) of positively stained cells by the staining intensity (0, negative; 1, weak; 2, moderate; and 3, strong). Scores greater than 6 indicated high expression, whereas scores less than 6 indicated low expression.

### Generation of knockout and stably over-expression cell lines

The CRISPR/Cas9 technique was used to generate gene knockouts. The annealed SET sgRNA oligonucleotides were cloned into the pLentiCRISPR-V2 vector. Human full-length SET was inserted into the pLVX-Myc-puro vector (by Age-I and EcoR-I).

SET knockdown or over-expression plasmids and virus packing plasmids were co-transfected into HEK293T cells to produce lenti-virus. The virus-containing supernatant was collected and purified after 48 hours, and BxPC3, Panc1, and Miapaca2 cells were then infected. The infected cells were selected with puromycin for 2 weeks. The genome sequence and western blot were used to identify all the knockout and over-expression cell lines.

### siRNA knockdown

RNA interference transfections were carried out in a forward transfection mode using Lipofectamine 3000 transfection reagent according to the manufacturer’s instructions. 1×10^5^ BxPC3 cells were seeded in 12-well plates before being transfected with a final siRNA mixture of 20 nM and 2 ul of Lipofectamine per ml. 72 hours after transfection, the cells were harvested for future experiments. Knockdown efficiency was examined by western blot.

### Western blot

Cells were lysed in 1×SDS-PAGE loading buffer containing DTT and incubated at 100°C for 10 minutes. β-Actin was utilized as a control for protein expression. All antibodies were determined experimentally and verified in the linear range. Samples were separated by SDS-PAGE (7.5–12.5%) and transferred onto the PVDF membranes. The membranes of PVDF were blocked with a TBS buffer containing 0.1% tween-20 and 5% non-fat milk or BSA. Then, the membranes were treated overnight at 4°C with a primary antibody diluted in the blocking buffer. The next day, membranes were incubated at room temperature with secondary antibodies conjugated to HRP for one hour. The membrane was finally constructed using a chemiluminescence substrate.

### RNA extraction and RT-qPCR

Total RNA was extracted from clinical tumor specimens and human PDAC cells using the RNA extraction kit. The cDNA was synthesized from total RNA using a Prime Script RT Reagent Kit according to the manufacturer’s instructions. The ΔΔCt technique was utilized to calculate relative mRNA expression levels, which were adjusted to β*2*M levels. ABI 7900HT Real-Time PCR machine (Applied Biosystems, Inc., USA) was used for thermal cycling. The primers used are listed in **Table S2**.

### DNA site-specific mutation

The site-specific mutagenesis in the SET or Pp2ac CDS was generated by the PCR-based method. The primers are listed in **Table S2**. Following the manufacturer’s instructions, the KOD-Plus mutagenesis kit (Toyobo, Osaka, Japan) was used to generate mutant plasmids. All truncated mutations were confirmed by Sanger sequencing and expression in the HEK293T cells.

### Cell viability assay

Cell viability was measured with the Cell Counting Kit-8 (CCK-8) following the manufacturer’s instructions. In the 96-well plates, 3000 cells were inoculated per well overnight. The next day, 10ul of CCK-8 solution were added to each well. After 90 minutes of incubation, the absorbance at 450 nm was measured with a microplate reader.

### Cell death-detecting drug efficacy assay

Control and SET-KO BxPC3 cells were seeded in the 12-well plates and added gradient tivantinib after cells adhered. 24 hours later, flow cytometric analysis was performed using the PE Annexin V Apoptosis Detection Kit according to the manufacturer’s instructions to determine cell apoptosis.

### Immunoprecipitation-mass spectrum (IP-MS)

1×10^7^ HEK293T and Panc02 cells were stably transfected with the control and Myc-SET plasmids, respectively. Magnetic beads conjugated with anti-SET antibody were pre-washed 5 times with 1× TBST buffer (25mM Tris, 0.15M NaCl, 0.05% Tween-20). Cells were washed three times with cold PBS and then lysed in cell lysis buffer containing an EDTA-free protease inhibitor cocktail. Add the cell samples to the pre-washed magnetic beads and incubate on a 4 °C roller overnight. Collect the beads with magnetic stand and wash the beads twice with 5× TBST buffer (125mM Tris, 0.75M NaCl, 0.25% Tween-20). Add 50 ul of SDS-PAGE loading buffer and incubate the sample at 100 °C for 10 minutes. The control SET and SET-OE associated proteins were immunoprecipitated with magnetic beads and subjected to SDS-PAGE separation on 10% gel.

Cut the protein gel strip into pieces and soak each piece for half an hour in sterile water in the corresponding EP tubes. Decolorize at 37°C for thirty minutes, or until the blue has faded. Add acetonitrile to dehydrate the gel and make it completely white twice. Soak up the acetonitrile and let it dry in the air. Add 10 mM DTT until the liquid covers the gel, then place in a 56 °C water bath for one hour. Place for 45 minutes in the dark after cooling to room temperature. Wash twice with the decolorizing solution. The enzyme was diluted with 25 mM NH_4_HCO_3_ and placed on ice for 30 minutes. Add the appropriate buffer to the gel and incubate at 37 °C overnight. Wash with 50% and 100% ACN and take the supernatant to be freeze-dried. Thermo UltiMate 3000 UHPLC was utilized to separate samples of dried peptides.

The liquid phase chromatography-separated peptides were ionized by a nanoESI source and passed to a tandem mass spectrometer Q-Exactive HF X for Data Dependent Acquisition mode detection.

### CUT&Tag

The CUT&Tag assay was performed with minor modifications as previously described (Kaya-Okur et al., 2019). Gently washed the cells three times with wash buffer and incubated them with activated concanavalin A magnetic beads at room temperature for 10 minutes. Add the primary antibody and incubate it on a rotating platform at 4°C overnight. Remove the primary antibody and incubate for 30 minutes at room temperature on a rotating platform with diluted secondary antibodies. Wash three times with Dig-Wash buffer. Add 0.04uM pA/G-Tnp and rotate incubation for 1 hour. The cells were resuspended in TruePrep tagging buffer and incubated for 1 hour at 37°C. Purified the DNA with DNA extract beads. Libraries were amplified using PCR according to the manufacturer’s protocol. Following the manufacturer’s recommendations, sequencing was carried out on an Illumina Novaseq 6000 utilizing 150 bp paired-end.

### Click-EU labeling nascent RNA

Cells were labeled with 5-ethynyluridine (5-EU) for 2 hours prior to fixation and permeabilization. Then wash the cells twice with 0.3% BSA in PBS and incubate at room temperature for 15 minutes with Click reaction buffer, CuSO4, Azide 488, and Click-IT additive solutons. Flow cytometry and imaging were performed on the treated cells.

### Nascent RNA sequencing

The nascent RNA sequencing was performed with minor modifications as previously described (Chen et al., 2018; Zheng et al., 2020). 1×10^7^ control and SET KO cells were harvested with trypsin and washed with ice-cold PBS for three times. Resuspend the cells in Hypotonic Buffer (10 mM pH = 7.9 HEPES; 10 mM KCl; 2 mM MgCl_2_; 1 mM DTT, 1× protease inhibitor cocktail) and incubate for 20 minutes on ice.

Centrifuge the suspension for 10 minutes at 4c and discard the supernatant. Use a pestle to pulverize the cells and ensure that 90% of the cells are broken up. Collect the nuclei by centrifuge at 600g for 10 minutes at 4c. Resuspend and wash the nuclei with Nuclei Wash Buffer (10mM pH 7.9 HEPES; 250mM Sucrose; 1mM DTT, 1×protease inhibitor cocktail; 50U/mL RNase inhibitor) for twice. Mix 50% volume of 2M fresh urea solution and 50% volume of cold 2× NUN Buffer (40 mM pH 7.9 HEPES; 15mM MgCl2; 600mM NaCl; 0.4mM EDTA; 2% v/v Nonidet P40) with DTT for 1×NUN Buffer. Suspend the pellet and break the nuclei by pipetting up and down 10–15 times with 1 mL of 1 NUN buffer. Incubate for 5 minutes at 4°C on a rotating plate. Spin the chromatin at 1000g for 3 minutes at 4°C to pelletize it. And carefully resuspend the chromatin pellet in 1×NUN buffer. Incubate the homogenized sample at 50°C for 5 minutes. Follow the standard RNA purification protocol with TRIzol for the remaining steps of RNA extraction.

Detected the nascent RNA with 1% agarose electrophoresis. Purge rRNA, synthesize cDNA, and construct and amplify libraries according to the manufacturer’s guidelines. Validate the quality of the library using the Agilent DNA 1000 Kit on the Agilent 2100 Bioanalyzer. It was suitable to have smooth profiles peaking around 250 bp and no detectable primer dimer contamination. After two days of sequencing on an Illumina Novaseq 6000 machine, the original data is finally converted into Fastq format.

### RNA-seq

RNA was isolated from control and SET-KO cells as previously described. On 0.8% agarose gels, RNA degradation and contamination were observed. The RNA sample preparations used a total of 3ug of RNA as input material for each sample. Utilizing the NEBNext® UltraTM RNA Library Prep Kit for Illumina® (NEB, USA) in accordance with the manufacturer’s instructions, sequencing libraries were created, and index codes were added to assign sequences to specific samples. Biological triplicates of samples were sequenced using the Illumina NovaSeq6000 platform. HTSeq v0.6.0 was utilized to determine the number of reads mapped to each gene. The TPM of each gene was then evaluated based on the gene’s length and the total number of mapped reads.

### ChIP-seq

Control and SET KO cells were harvested for ChIP assay utilizing validated ac-H4 or Pol II (Rpb1) NTD antibodies. Each group underwent two biological replicates of experiments. ChIP was performed based on the manufacturer’s instructions using SimpleChIP® Plus Enzymatic Chromatin IP Kit (Magnetic Beads). Specifically, approximately 5×10^6^ cells were first fixed with 1% formaldehyde fixation. In each spike-in ChIP experiment, human cells were utilized in a 50:1 ratio to Drosophila S2 cells. At the start of the ChIP workflow, S2 cells were mixed in with human cells.

After combining Drosophila S2 and human cells, the sample was treated as one ChIP-seq sample all across the experiment until the DNA sequencing was completed as described previously (Wu et al., 2021a; Wu et al., 2021b). The input and DNA samples were used to prepare sequencing libraries in accordance with the Illumina Genome Analyzer manufacturer’s recommendations. After amplification, the total amount of DNA in each specimen was standardized. An Illumina novaseq 6000 Analyzer was used to sequence the purified.

### PP2A immunoprecipitation phosphatase assay

For PP2A phosphatase assay, immunoprecipitations using lysates from 6-well plate were performed according to the manufacturer’s instructions. Briefly, wash the cells with 1× TBS for three times, and scrape the plates with 300ul phosphatase extraction buffer (20mM imidazole-HCl, 2mM EGTA, 2mM EDTA, 1mM benzamidine, 1×PMSF, pH 7.0, 1×protease inhibitor cocktail). Sonicate the cells for 10 seconds and centrifuge for 10minutes at 2000g at 4c. The tests for phosphatase activity were conducted using the isolated supernatant. Add PP2A-C antibody, Protein A agarose, and pNPP Ser/Thr Assay buffer, and then incubate for 1 hour at 4°C on constantly rocking plate. Wash and incubate with Ser/Thr Assay Buffer for 8 minutes at 30c with shaking. Add malachite green phosphate detection solution and read at 650nm.

## QUANTIFICATION AND STATISTICAL ANALYSIS

All replicate experiments were conducted using a minimum of two biological replicates per condition. Statistical tests were performed after confirming that the data met appropriate assumptions (ex. Normal distribution) and two-tailed. P-values were calculated and adjusted for multiple hypothesis testing where indicated. Statistical significance is indicated as *p < 0.05, ** p < 0.01, *** p < 0.001. The statistical analyses were performed with R and GraphPad Prism. Pearson χ2 test or Fisher’s exact test was used to examine categorical variables. Student’s t-test and Mann-Whitney-Wilcoxon test were used to analyze continuous variables. Survival comparisons were evaluated using Kaplan-Meier curves and log-rank tests. In RT-qPCR, relative gene expression △Ct (critical threshold) = Ct of genes of interest-Ct of β*2*M. In ChIP-qPCR, the enrichment for specific locus was normalized by input in the same way. Fold changes of gene expression level or ChIP enrichment were calculated by the 2^-Ct^ method. The data in histograms were presented as the mean±standard deviation.

### Cell transcriptome connectivity matching

The Cell Connectivity Map (CMAP) is a compendium of gene expression signatures induced by chemical compounds or genetic perturbations (termed perturbagens) in multiple cell lines. For each small molecule treatment or single gene knockdown/over-expression in a specific cell line, the L1000 assay platform measured the reduced representation of total transcriptome (∼1,000 land mark transcripts), recorded as an expression signature (Lamb et al., 2006; Subramanian et al., 2017).

CMAP has collected approximately 1.3 million expression signatures of over 8,000 perturbagens in at least 7 cell types, and the “distance” between any two expression signatures is quantified using Kolmogorov-Smirnov test. More similar expression signatures showed “shorter distance” and related higher “connectivity value”., We searched CMAP (https://clue.io/) using the command: /conn “CDK9-OE”. The resultant connectivity data was then downloaded and visualized as heatmaps.

### Nascent RNA-seq analysis

Raw reads were trimmed and then aligned to human hg19 and drosophila dm6 genome assemblies using STAR v2.7.5c with parameter ‘‘–outSAMtype BAM SortedByCoordinate–twopassMode Basic–outFilterMismatchNmax 2–outSJfilterReads Unique’’ (Dobin et al., 2013). Read normalization was based on 1e6/spike-in read number calculated by SAMtools v1.9 and normalized bigwig files were built with deeptools v3.5.0 (Li et al., 2009; Ramirez et al., 2016). Gene expression quantification was performed with featureCounts tool from the Rsubread R package v2.0.1 (Liao et al., 2019). For each gene, we computed the number of fragments per kilobase of exon per million mapped reads (FPKM) by normalizing the spike-in dm6 reads. Differential expression analysis was performed using DESeq2 R package v1.26.0 (Love et al., 2014).

### ChIP-seq analysis

The paired-end ChIP-seq reads were processed with Trim Galore v0.6.6 (https://www.bioinformatics.babraham.ac.uk/projects/trim_galore/) to remove adaptors and low-quality reads with the parameter ‘‘-q 25’’ and then aligned to the human hg19 and drosophila dm6 assemblies using Bowtie v2.3.5.1 with default parameters (Langmead and Salzberg, 2012). All unmapped reads, low mapping quality reads (MAPQ < 30) and PCR duplicates were removed using SAMtools v1.9 (Li et al., 2009) and Picard v2.23.3 (https://broadinstitute.github.io/picard/). The number of spike-in dm6 reads was counted with SAMtools v1.9 and normalization factor alpha = 1e6/dm6_count was calculated (Li et al., 2009). Normalized bigwig was generated with deeptools v3.5.0 and reads mapped to the ENCODE blacklist regions were removed using bedtools v2.29.2 (Amemiya et al., 2019; Quinlan and Hall, 2010; Ramirez et al., 2016). Peaks were called using macs2 v2.2.6 with a q-value threshold of 0.05 and peak annotation was performed with ChIPseeker R package v1.24.0 (Yu et al., 2015; Zhang et al., 2008).

### CUT&Tag analysis

Raw CUT&Tag reads were processed. CUT&Tag reads were trimmed using cutadapt (v1.16) (-m 10 -q 10). Paired-end reads were aligned to the hg19 reference genome using bowtie2 with default parameters. Normalized bigWig files were generated for each replicate using the bamCoverage function in the deeptools package. Replicates were further normalized to the input using the bigwigCompare function with the parameter –operation log.

### Gene ontology

Gene ontology (GO) analysis was performed using the web interface for Database for Annotation, Visualization, and Integrated Discovery (DAVID, v.6.8). The Functional Annotation tool was used to identify biological process GO terms (GOTERM_BP_DIRECT) that are enriched in the queried gene list. Selected GO terms with Benjamini-Hochberg adjusted p-value < 0.05 were plotted.

### Data visualization

Reads per genomic content (RPGC)-normalized bigWig files were generated from ChIP-seq, CUT&Tag, nascent RNA-seq and RNA-seq bam files using the BamCoverage function in the deeptools package (v.3.0.1) with parameters –normalizeUsing RPGC –effectiveGenomeSize 2652783500 –centerReads. For ChIP-seq and CUT&Tag data, the parameter –extendReads 150 was also used. RPGC-normalized ChIP-seq, CUT&Tag, nascent RNA-seq and RNA-seq tracks at specific loci were visualized using Integrative Genomics Viewer (IGV, Broad Institute).

RPGC-normalized ChIP-seq and CUT&Tag signal was calculated in regions ±3kb centered at the enhancer using the computeMatrix function in the deeptools package (reference-point –referencePoint center -a 3000, -b 3000 –binSize 10). Heatmaps and profiles of the signal were visualized using plotHeatmap and plotProfile functions respectively using default parameters (**Figures 2C, 5G and S2B**). Custom R code was used to generate profiles of mean signal from different samples at the same set of regions (**Figures 2E**, **5G, 5H, S5C and S5E)**.

**Table S1** summarized the relationship between the sequencing data and the derived figure results.

---

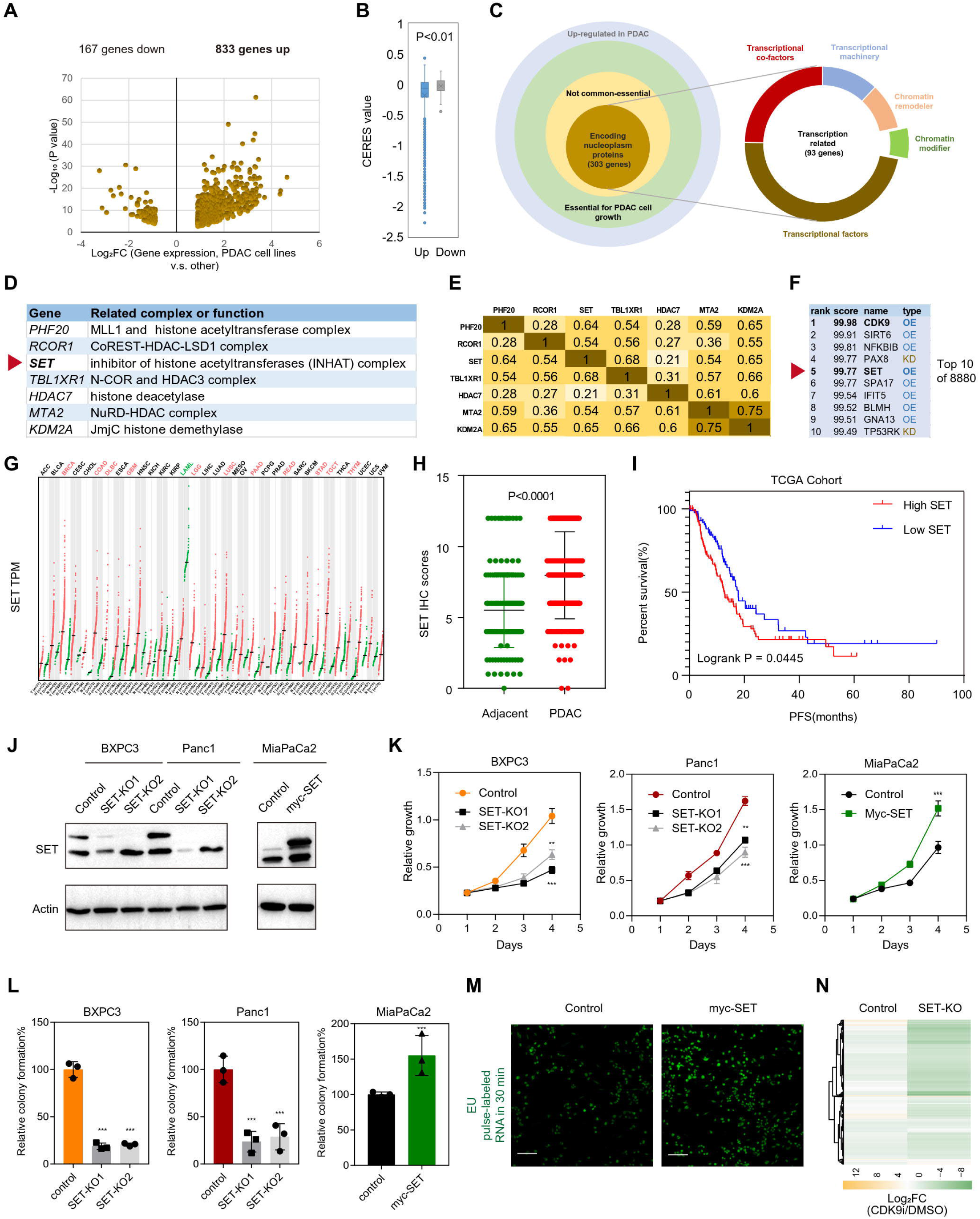
Figure S1.

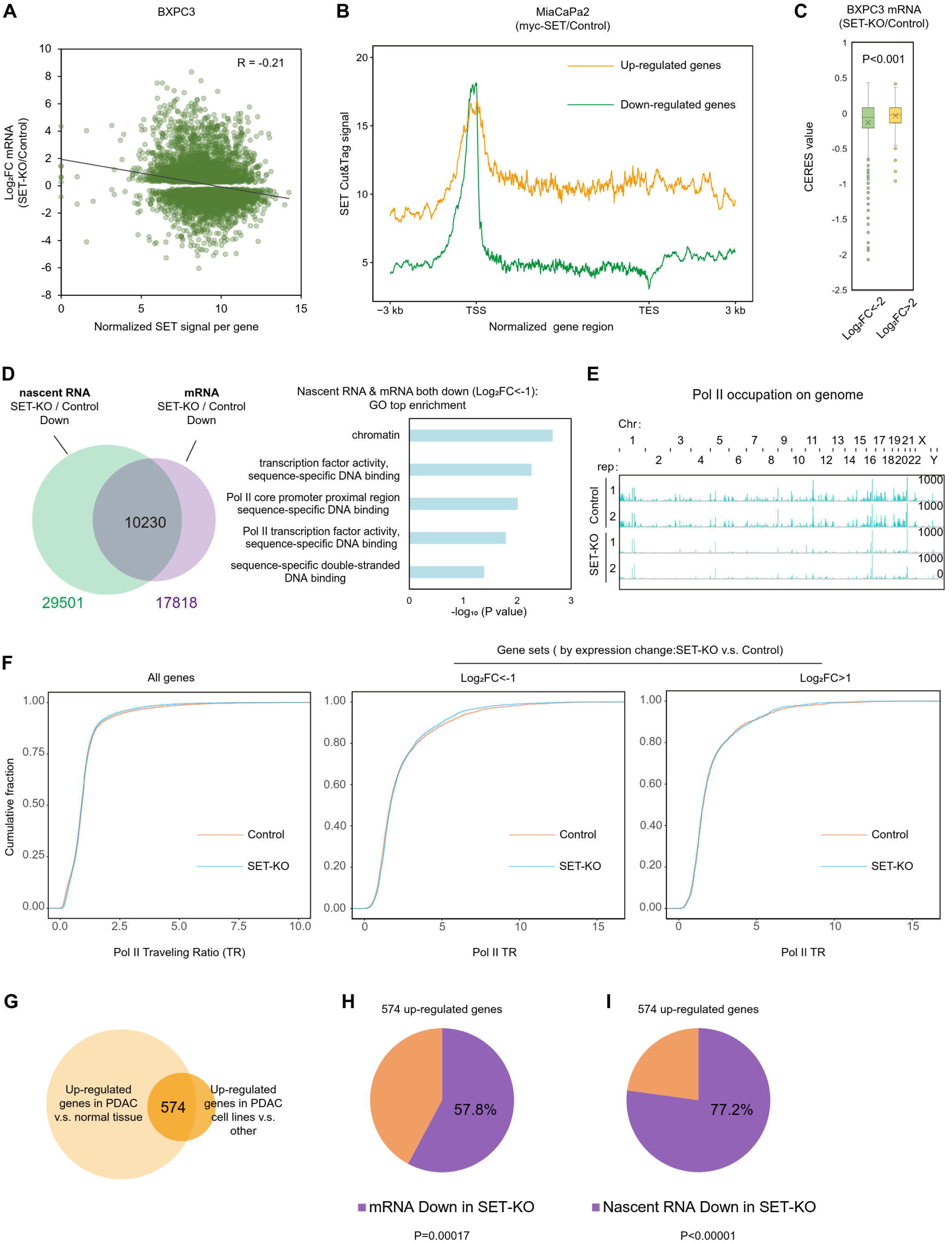
Figure S2.

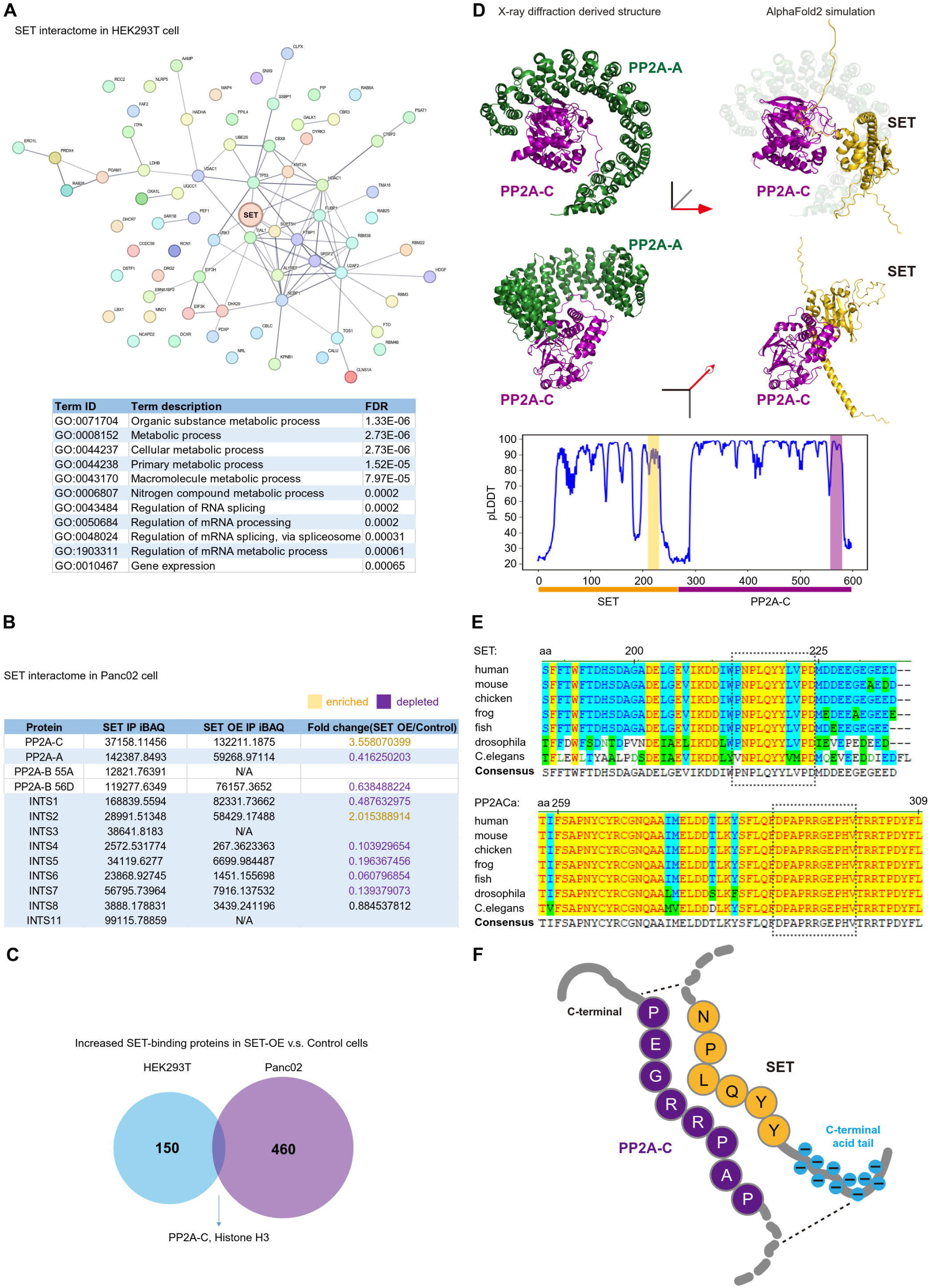
Figure S3.

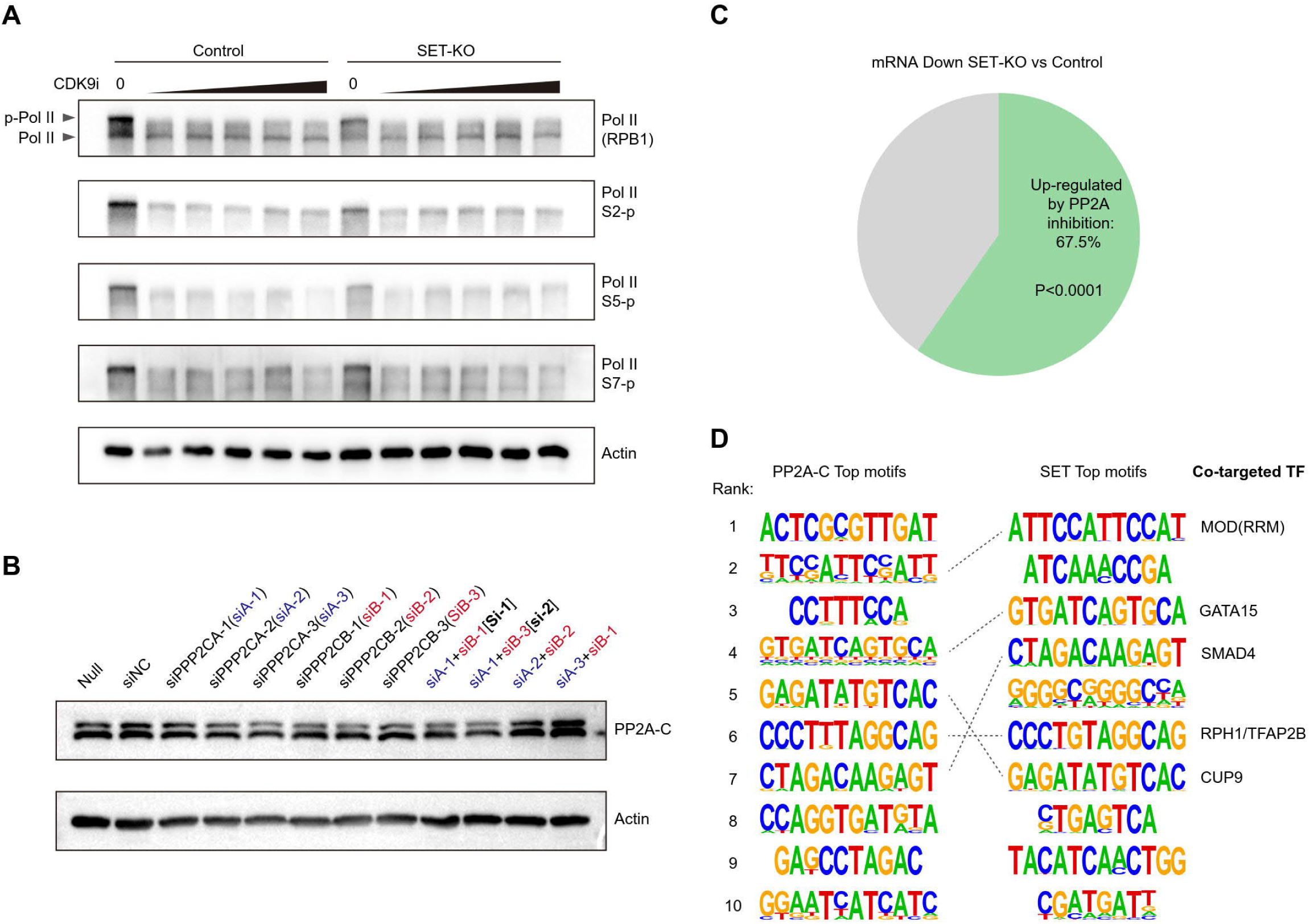
Figure S4.

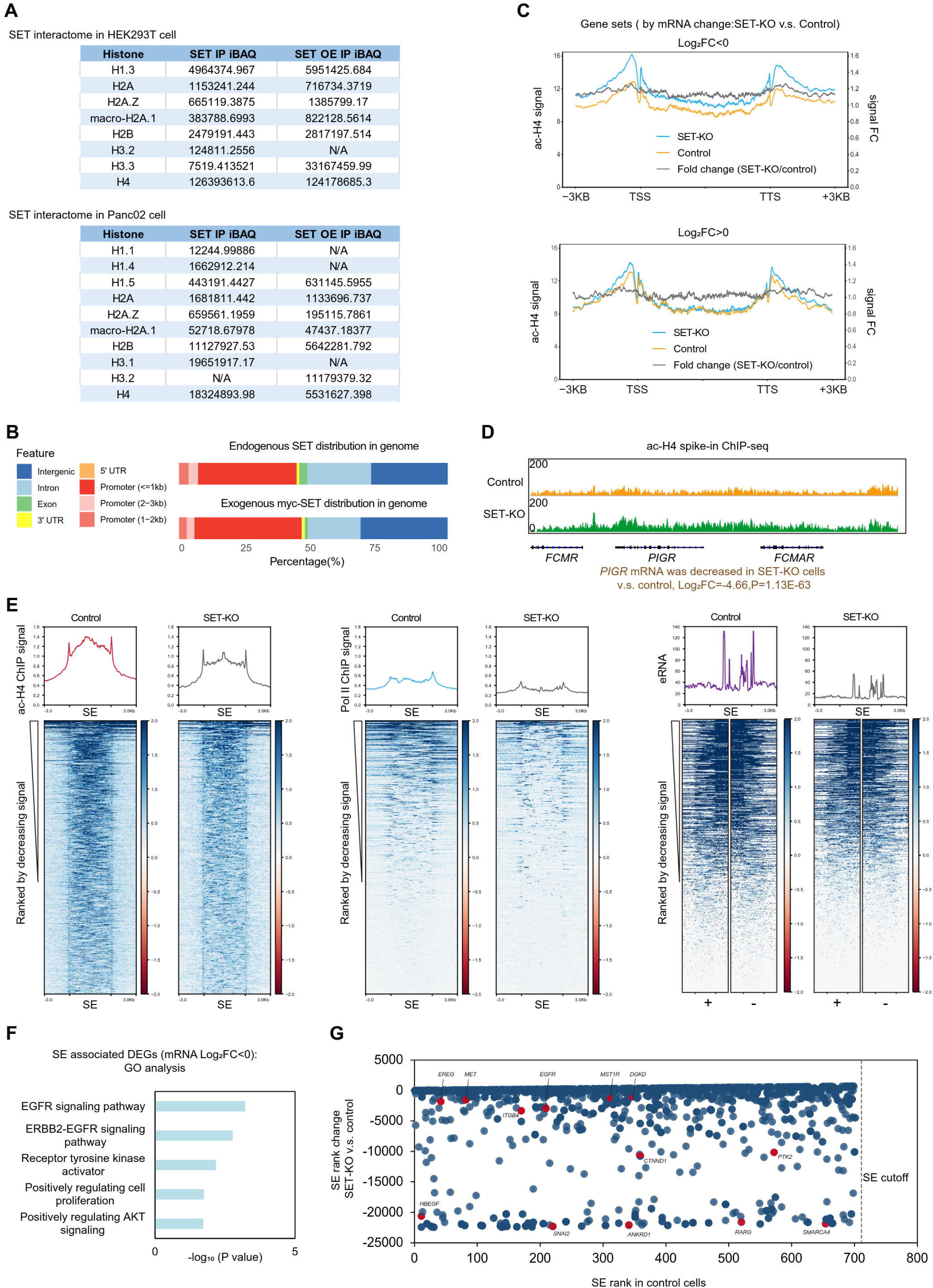
Figure S5.

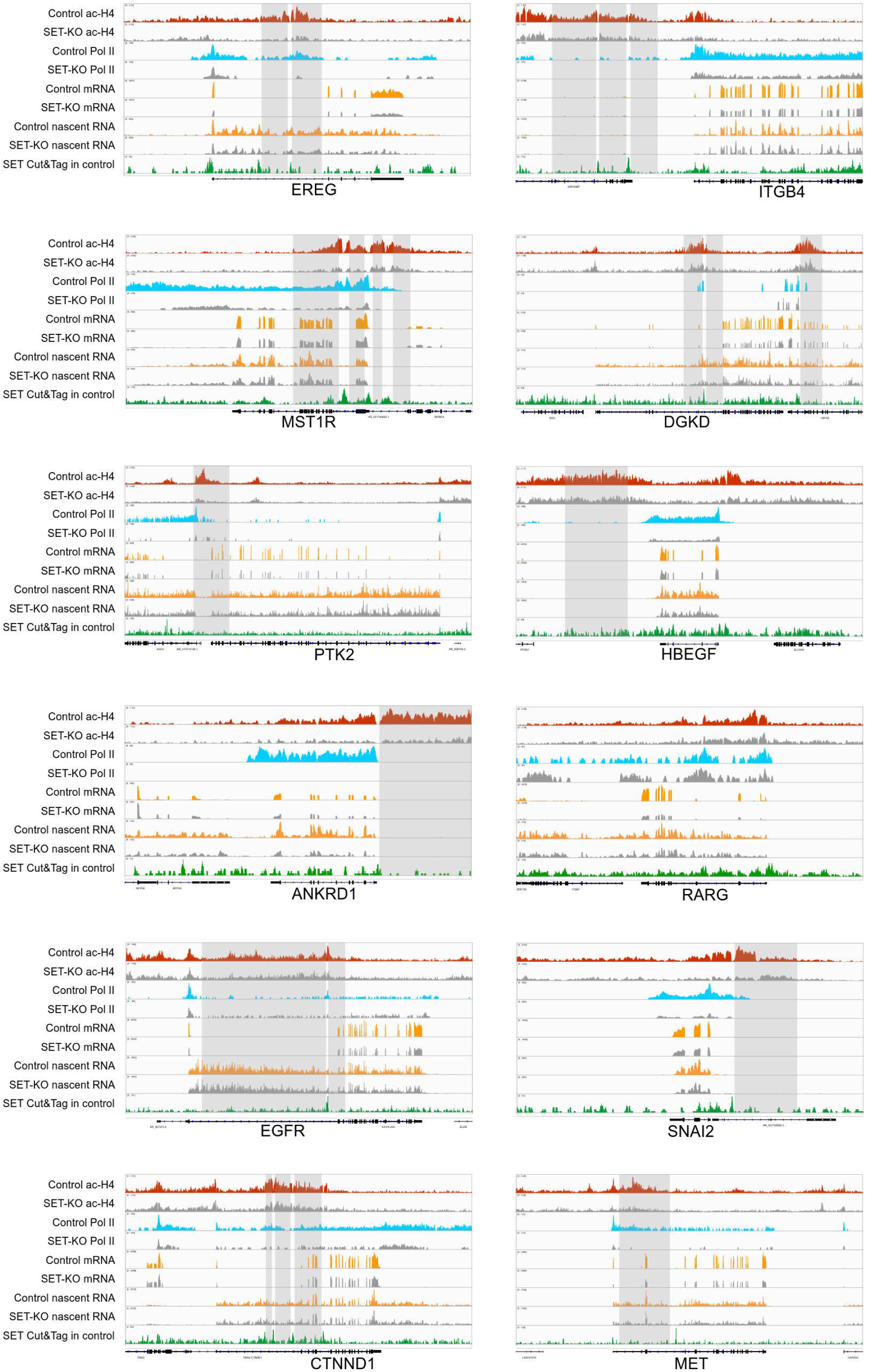
Figure S6.

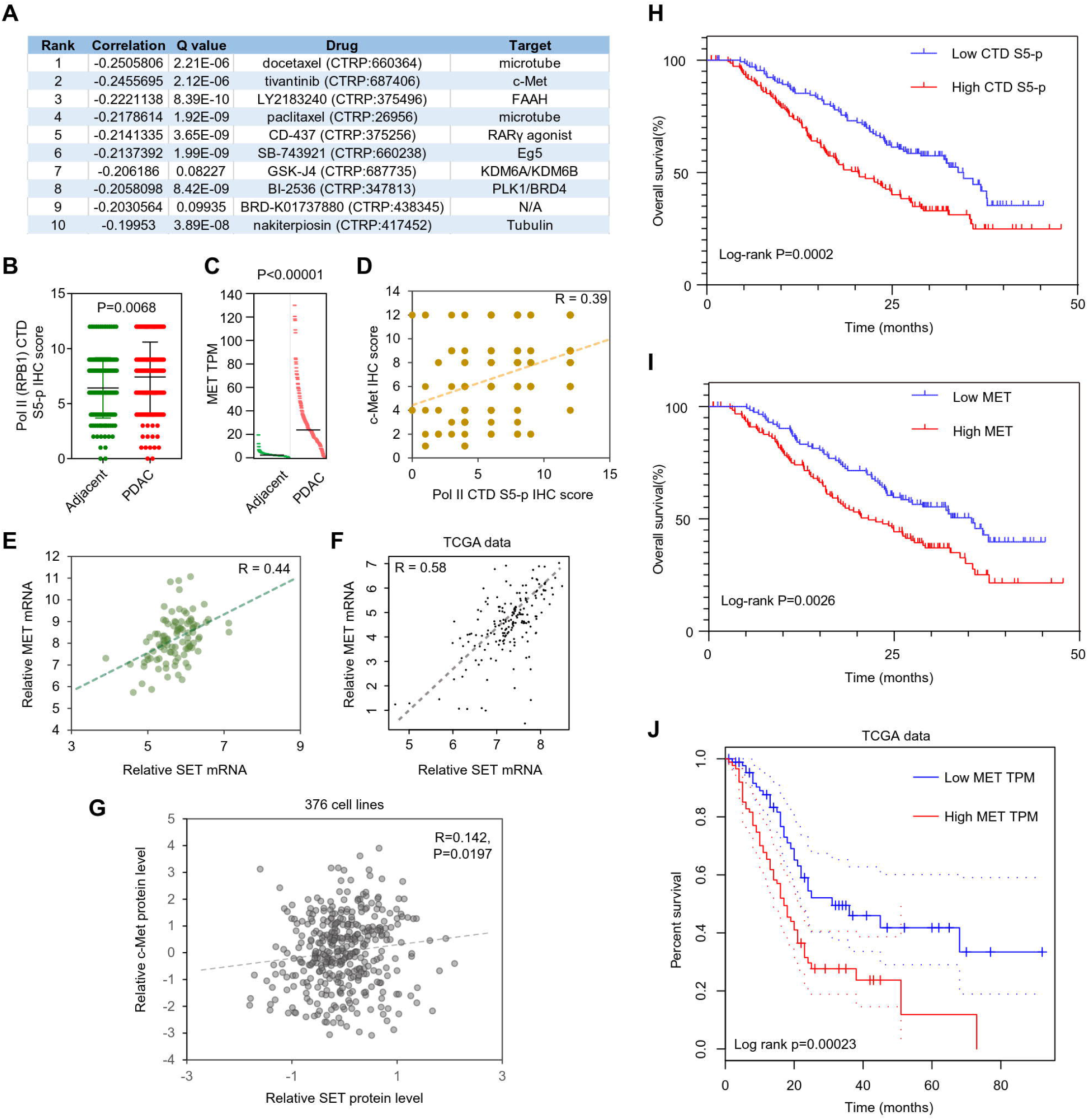
Figure S7.

