## Supplementary material for "Histone acetylation insulator SET orchestrates PP2A inhibition and super-enhancer activation": Table S1

**Table S1. Navigation to the relationship between the raw sequencing data and the derived figure results.**

| **Figure** | **Related NGS type** | **Related NGS track** |
| --- | --- | --- |
| Figures 1I-1M | Nascent RNA-seq | Control+dmso.bw |
|  |  | Control+CDK9i.bw |
|  |  | SET-KO+dmso.bw |
|  |  | SET-KO+CDK9i.bw |
| Figures 2A, 2B | RNA-seq | BXPC3-Control.bw |
|  |  | BXPC3-SET-KO.bw |
| Figure 2C | RNA-seq, Cut & Tag | BXPC3-Control.bw |
|  |  | BXPC3-SET-KO.bw |
|  |  | BXPC-SET-rep1.bw |
|  |  | BXPC-SET-rep2.bw |
| Figure 2E | ChIP-seq | Control_Pol II_rep1.bw |
|  |  | Control_Pol II_rep2.bw |
|  |  | SET_KO_Pol II_rep1.bw |
|  |  | SET_KO_Pol II_rep2.bw |
| Figure 4E, 4I and 4J | RNA-seq | BXPC3 control or SET-KO treated with LB100 (RNA-seq gene TPM list).xlsx |
| Figure 4G | Cut & Tag | BXPC-PP2AC-rep1.bw |
|  |  | BXPC-PP2AC-rep2.bw |
|  |  | BXPC-SET-rep1.bw |
|  |  | BXPC-SET-rep2.bw |
| Figures 5E-5J | ChIP-seq, Cut & Tag | Control_acH4_rep1.bw |
|  |  | Control_acH4_rep2.bw |
|  |  | SET_KO_acH4_rep1.bw |
|  |  | SET_KO_acH4_rep2.bw |
|  |  | Control_Pol II_rep1.bw |
|  |  | Control_Pol II_rep2.bw |
|  |  | SET_KO_Pol II_rep1.bw |
|  |  | SET_KO_Pol II_rep2.bw |
|  |  | BXPC-PP2AC-rep1.bw |
|  |  | BXPC-PP2AC-rep2.bw |
|  |  | BXPC-SET-rep1.bw |
|  |  | BXPC-SET-rep2.bw |
| Figure S2A | RNA-seq, Cut & Tag | BXPC3-Control.bw |
|  |  | BXPC3-SET-KO.bw |
|  |  | BXPC-SET-rep1.bw |
|  |  | BXPC-SET-rep2.bw |
| Figure S2B | RNA-seq, Cut & Tag | Mia-Control.bw |
|  |  | Mia-SET_OE.bw |
|  |  | Mia-myc-SET.bw |
| Figure S2D | RNA-seq, Nascent RNA-seq | Control+dmso.bw |
|  |  | SET-KO+dmso.bw |
|  |  | BXPC3-Control.bw |
|  |  | BXPC3-SET-KO.bw |
| Figures S2E, S2F | ChIP-seq | Control_Pol II_rep1.bw |
|  |  | Control_Pol II_rep2.bw |
|  |  | SET_KO_Pol II_rep1.bw |
|  |  | SET_KO_Pol II_rep2.bw |
| Figures S2H, S2I | RNA-seq | BXPC3-Control.bw |
|  |  | BXPC3-SET-KO.bw |
| Figure S4C | RNA-seq | BXPC3-Control.bw |
|  |  | BXPC3-SET-KO.bw |
|  |  | BXPC3 control or SET-KO treated with LB100 (RNA-seq gene TPM list).xlsx |
| Figure S4D | Cut & Tag | BXPC-PP2AC-rep1.bw |
|  |  | BXPC-PP2AC-rep2.bw |
|  |  | BXPC-SET-rep1.bw |
|  |  | BXPC-SET-rep2.bw |
| Figure S5B | Cut & Tag | BXPC-SET-rep1.bw |
|  |  | BXPC-SET-rep2.bw |
|  |  | Mia-myc-SET.bw |
| Figures S5C-S5G | ChIP-seq, Nascent RNA-seq, RNA-seq | Control_acH4_rep1.bw |
|  |  | Control_acH4_rep2.bw |
|  |  | SET_KO_acH4_rep1.bw |
|  |  | SET_KO_acH4_rep2.bw |
|  |  | Control_Pol II_rep1.bw |
|  |  | Control_Pol II_rep2.bw |
|  |  | SET_KO_Pol II_rep1.bw |
|  |  | SET_KO_Pol II_rep2.bw |
|  |  | Control+dmso.bw |
|  |  | SET-KO+dmso.bw |
|  |  | BXPC3-Control.bw |
|  |  | BXPC3-SET-KO.bw |
