## Supplementary material for "Histone acetylation insulator SET orchestrates PP2A inhibition and super-enhancer activation": Table S2

**Table S2. oligonucleotide sequences**

| **sgRNA target sequences** | |
| --- | --- |
| sgSET-1 | GCCTAGTGTCTGCACTGCTT |
| sgSET-2 | GAGACTGCCCCAGGTGCCAG |
| **siRNAs** | |
| siPPP2CA-1 | ACCGGAAUGUAGUAACGAUUU |
| siPPP2CA-2 | GGCAAAUCACCAGAUACAAAU |
| siPPP2CA-3 | CCCAUGUUGUUCUUUGUUAUU |
| siPPP2CB-1 | AGCCGACAAAUUACCCAAGUA |
| siPPP2CB-2 | UGUCUGCGAAAGUAUGGGAAU |
| siPPP2CB-3 | GAAAUCACGAAAGCCGACAAA |
| **qPCR primers** | |
| SET-F | AGCAAGAAGCGATTGAACACA |
| SET-R | TGGTTGGCGGAGTTTGTTATATT |
| MET-F | AGCAATGGGGAGTGTAAAGAGG |
| MET-R | CCCAGTCTTGTACTCAGCAAC |
| EGFR-F | TTGCCGCAAAGTGTGTAACG |
| EGFR-R | GTCACCCCTAAATGCCACCG |
| ZBTB5-F | CCATTTCAGATGTTACACCGGA |
| ZBTB5-R | TCGCACTATCTTCCTGGTTATCA |
| **Mutation primers** | |
| SET-del F | TAAGAATTCGCCAATTCGCTGGAGC |
| SET-del C1 R | CATATCGGGAACCAAGTAGTACTGT |
| SET-del C2 R | ACCTGCATCAGAATGGTCAGTAAAC |
| SET-del C3 R | TTCGGTGGACTTCGAAGATGGATCA |
| SET-del C4 R | GCTGGCTTTATTCTGCGTTTGACTC |
| Pp2ac-del N F | AAGGTTCGTTACCGAGAGCGCATCAC |
| Pp2ac-del N R | CATGGTGGCGGGATCCATTCCACTAC |
| Pp2ac-del M F | GGTGGCTGGGGGATATCTCCTCGGG |
| Pp2ac-del M R | AAGAGCTACAAGCAGTGTAACTGTTTC |
| Pp2ac-del C F | GATCCCGCCACCATGGTGAGCAAGG |
| Pp2ac-del C F | ACGGTCATCTGGATCTGACCACAGC |
